# Characterization of SARS2 Nsp15 Nuclease Activity Reveals it’s Mad About U

**DOI:** 10.1101/2021.06.01.446181

**Authors:** Meredith N. Frazier, Lucas B. Dillard, Juno M. Krahn, Lalith Perera, Jason G. Williams, Isha M. Wilson, Zachary D. Stewart, Monica C. Pillon, Leesa J. Deterding, Mario J. Borgnia, Robin E. Stanley

**Affiliations:** Signal Transduction Laboratory, National Institute of Environmental Health Sciences, National Institutes of Health, Department of Health and Human Services, 111 T. W. Alexander Drive, Research Triangle Park, NC 27709, USA; Genome Integrity and Structural Biology Laboratory, National Institute of Environmental Health Sciences, National Institutes of Health, Department of Health and Human Services, 111 T. W. Alexander Drive, Research Triangle Park, NC 27709, USA; Epigenetics and Stem Cell Biology Laboratory, National Institute of Environmental Health Sciences, National Institutes of Health, Department of Health and Human Services, 111 T. W. Alexander Drive, Research Triangle Park, NC 27709, USA

## Abstract

Nsp15 is a uridine specific endoribonuclease that coronaviruses employ to cleave viral RNA and evade host immune defense systems. Previous structures of Nsp15 from across *Coronaviridae* revealed that Nsp15 assembles into a homo-hexamer and has a conserved active site similar to RNase A. Beyond a preference for cleaving RNA 3’ of uridines, it is unknown if Nsp15 has any additional substrate preferences. Here we used cryo-EM to capture structures of Nsp15 bound to RNA in pre- and post-cleavage states. The structures along with molecular dynamics and biochemical assays revealed critical residues involved in substrate specificity, nuclease activity, and oligomerization. Moreover, we determined how the sequence of the RNA substrate dictates cleavage and found that outside of polyU tracts, Nsp15 has a strong preference for purines 3’ of the cleaved uridine. This work advances our understanding of how Nsp15 recognizes and processes viral RNA and will aid in the development of new anti-viral therapeutics.

## INTRODUCTION

The novel SARS-CoV-2 (severe acute respiratory syndrome coronavirus 2) emerged in late 2019 and became a worldwide pandemic that is still ongoing and has infected millions worldwide (1). Coronaviruses are members of the Nidovirus order, which encompasses large, positive-strand RNA viruses with genomes that range in size from 12-41 kb (2). The 30 kb SARS-CoV-2 genome encodes for 4 structural proteins that are part of the mature viral particle, 8 accessory proteins, and 15 non-structural proteins (Nsps) (3). The Nsps are encoded in two open reading frames found in the first two-thirds of the viral genome. These proteins are translated by host ribosomes as two long polyproteins and are cleaved into functional proteins by the viral proteases (4). The Nsps play important roles in viral replication and pathogenicity and many of them are promising drug targets (3,4).

Nsp15 is a uridine specific endoribonuclease conserved across the *Coronaviridae* family (5). Enzymatic activity occurs in the C-terminal EndoU domain, which is more broadly conserved across nidoviruses, suggesting that this endoribonuclease activity is critically important for large, positive-strand RNA viruses (5,6). Work in animals and cell culture has shown that Nsp15 function is not necessary for viral replication, however Nsp15 nuclease activity is critically important for evasion of the host immune response to the virus, specifically by preventing the activation of dsRNA sensors (7–11). For example, in studies of porcine endemic diarrhea virus (PEDV), Nsp15-deficient virus resulted in higher levels of type I and III interferon responses in cells, and piglets infected with the mutant virus had much higher survival rates than those infected with WT PEDV (8). A similar effect was also seen in studies of mouse hepatitis virus (MHV); mice immunized with Nsp15 nuclease deficient virus were able to successfully clear WT virus, with commonly affected organs showing no pathology (7). Recent work also revealed a similar trend in the chicken infection bronchitis virus (IBV), where animals infected with nuclease-deficient virus had reduced mortality and viral shedding (12). Therefore, Nsp15 is a promising therapeutic target for coronaviruses.

One of the major outstanding questions about the function of Nsp15 is what is its RNA target for cleavage (5). Recent studies have begun to shed light on this important question (9,13). One study found a link between Nsp15 activity and the length of the polyuridine sequence at the 5’ end of the template negative strand. When Nsp15-mutant MHV infected cells, there was a greater amount of polyuridine (PUN) RNA compared to cells infected with WT virus, suggesting Nsp15 cleaves the PUN RNA produced in the negative strand intermediate state (9). The PUN RNA was found to trigger the pathogen associated molecular pattern (PAMP) receptor MDA5, which mediates interferon response (9). Another study used cyclic phosphate RNA sequencing to identify Nsp15 cleavage products within MHV infected bone marrow-derived macrophages (13). This analysis revealed that Nsp15 cleaves numerous targets throughout the positive strand with a preference for cleaving between U^A and C^A sequences (13). More recent work with IBV demonstrated that Nsp15 nuclease activity prevents the accumulation of both dsRNA and cytoplasmic stress granules which have established anti-viral properties (14). Collectively these studies confirm that Nsp15 nuclease activity is critical to prevent the accumulation of viral ds-RNA and activation of the immune response.

Numerous structures of Nsp15 have been determined from several *Coronaviridae* family members, however there are no structures of Nsp15 with more than a di-nucleotide bound, which has hindered our understanding of how Nsp15 recognizes its RNA targets. Crystal structures of Nsp15 revealed that Nsp15 assembles into a hexameric complex, formed from back-to-back trimers with the EndoU domains facing outward (15–19). The active site of Nsp15 shares considerable similarity to the well-studied endoribonuclease RNase A, and is composed of a catalytic triad including two histidines and lysine (18,20,21). These residues support a two-step reaction of transesterification and hydrolysis, however due to an altered position of one active site histidine, Nsp15 accumulates products from the transesterification reaction which contain a cyclic phosphate (20). Recent structures determined of Nsp15 bound to uridine nucleotides uncovered the molecular basis for uridine specificity, which is driven by a well-conserved serine residue within the uridine binding pocket (20,21). However, beyond the preference for uridines, it is unclear if Nsp15 has any additional specificity requirements. RNA sequencing suggests there is a preference for adenine 3’ to the uridine, but the structural basis for this is unknown (13). In contrast, RNase A is known to have additional non-catalytic sites that affect substrate preference (22,23), prompting further characterization of how Nsp15 engages RNA.

Here we used cryo-EM, molecular dynamics simulations, and in vitro RNA cleavage assays to probe the substrate specificity of SARS-CoV-2 Nsp15. We determined cryo-EM reconstructions of Nsp15 with RNA bound in the pre- and post-cleavage states. The structures revealed that, in contrast to RNase A, Nsp15 does not contain any additional well-ordered sites for RNA binding and recognition. This observation was further supported by molecular dynamics simulations with tri-nucleotide substrates. We probed RNA specificity by determining how the nucleotide 5’ and 3’ of the uridine affects cleavage and found that Nsp15 has a preference for purines 3’ of the cleaved pyrimidine. Finally, we looked at Nsp15’s ability to cleave SARS-CoV-2 viral RNA substrates, such as the PUN and the transcriptional regulatory sequence (TRS). Collectively our work suggests that SARS CoV-2 Nsp15 is able to cleave a broad spectrum of RNA substrates and that this activity is driven by recognition of uridine within the active site.

## MATERIAL AND METHODS

### Protein expression and purification

Wild type (WT) and mutant Nsp15 constructs were created as described previously (20). Nsp15 was overexpressed in *E. coli* C41 (DE3) competent cells in Terrific Broth with 100 mg/L ampicillin. At an optical density (600 nm) between 0.8-1.0, cultures were cooled at 4°C for 1 hour prior to induction with 0.2 mM Isopropyl β-D-1-thiogalactopyranoside (IPTG). Cells were harvested after overnight expression at 16°C and stored at −80°C until use. Nsp15 purification was done as described previously (20). Briefly, cells were resuspended in Lysis Buffer (50 mM Tris pH 8.0, 500 mM NaCl, 5% glycerol, 5 mM β-ME, 5 mM imidazole) supplemented with complete EDTA-free protease inhibitor tablets (Roche) and disrupted by sonication. The lysate was clarified at 26,915 x g for 50 minutes at 4°C and then incubated with TALON metal affinity resin (Clontech). His-Nsp15 was eluted from the resin with 250 mM imidazole, and buffer exchanged into Thrombin Cleavage Buffer (50 mM Tris pH 8.0, 150 mM NaCl, 5% glycerol, 2 mM β-ME, 2 mM CaCl_2_) for cleavage at room temperature for 3 hours. The cleavage reaction was repassed over TALON resin and quenched with 1 mM phenylmethylsulfonyl fluoride (PMSF) prior to gel filtration using a Superdex-200 column equilibrated in SEC buffer (20 mM Hepes pH 7.5, 150 mM NaCl, 5 mM MnCl_2_, 5 mM β-ME).

### Cryo-EM sample preparation

Purified Nsp15 was diluted in a low-salt buffer (20 mM Hepes pH 7.5, 100 mM NaCl, 5 mM MnCl_2_, 5 mM β-ME) to 0.75 μM and incubated with excess RNA substrates (1 mM AU^f^A or AUA, see Table S1) for 1 hour at 4°C. UltrAuFoil R1.2/1.3 300 mesh gold grids (Quantifoil) were plasma cleaned (Pie Scientific) before use. The Nsp15/RNA mixture (3 μL) was deposited onto the grids, back-blotted for 3 seconds, and vitrified using an Automatic Plunge Freezer (Leica).

### Data collection and processing

Nsp15 images were collected using a Krios electron microscope at 300 keV with a Gatan K2 detector in super-resolution mode. Beam-induced motion and drift were corrected using MotionCor2 (24) and aligned dose-weighted images were used to calculate CTF parameters using CTFFIND4 (25). CryoSPARC v2 (26) was used in all subsequent image processing. Particles were selected by template-based particle picking, downsampled by a factor of 4, extracted with a box size of 64 and subjected to an initial round of 2D classification. Full resolution particle projections from good classes were re-extracted using a box size of 256. Ab initio reconstruction was used to generate an initial model. Three independent 3D refinement cycles were performed while applying C1, C3, and D3 symmetry respectively. Although previous apo and UTP-bound datasets had D3 symmetry, the longer RNA bound in both datasets here resulted in particles that no longer had D3 symmetry, perhaps due to incomplete or mixed occupancy. Inspection of the C1 map did not reveal any asymmetric differences, although active site density was difficult to interpret for one half of the pre-cleavage state map. Therefore, C3 symmetry was used for model building and analysis for both datasets. Maps were re-scaled to optimize RMS fit to core domain residues of reference structure PDBID 6WLC (21).

### Model building

A SARS-CoV-2 Nsp15 crystal structure (PDBID 6WLC) was used as a starting model and fit into the cryo-EM maps using rigid body docking in Phenix (27). For the pre-cleavage state, which was captured with an AU^f^A tri-nucleotide, the density for the 5’ A was weaker than the density for the U, so only the C5’ group was modeled; no density was observed for the 3’ A. For the post-cleavage state, the 5’ A could be fit in the density along with the U. A combination of rigid body and real-space refinement in Phenix as well as iterative rounds of building in COOT (28) were used to improve the fit of the model. Molprobity (29) was used to evaluate the model (Table 1). Figures were prepared using Chimera (30) and Chimera X (31).

**Table 1.** Cryo-EM collection and processing statistics for the pre-cleavage and post-cleavage structures.

| **Data collection and processing** | Pre-cleavage state<br>EMBD-24137<br>PDB ID:7N33 | Post-cleavage state<br>EMBD-24101<br>PDB ID: 7N06 |
| --- | --- | --- |
| Microscope | Titan Krios | Titan Krios |
| Detector | Gatan K2 | Gatan K2 |
| Nominal magnification | 165000x | 165000x |
| Voltage (kV) | 300 | 300 |
| Electron exposure (e<sup>-</sup>/Å<sup>2</sup>) | 54 | 54 |
| Defocus range (μm) | -0.8 to -1.8 | -0.8 to -1.8 |
| Pixel size (Å) | 0.4125 | 0.4125 |
| Symmetry imposed | C3 | C3 |
| Number of Micrographs | 2563 | 3454 |
| Initial particle images | 944,302 | 1,592,506 |
| Final particle images | 383,275 | 1,058,228 |
| Map resolution (Å)/FSC threshold | 2.5/0.143 | 2.2/0.143 |
| **Refinement** |  |  |
| Resolution (Å) | 2.5 | 2.2 |
| B-factor used for map sharpening (Å<sup>2</sup>) | 115.5 | 94.5 |
| **Map to model CC** |  |  |
| CC (mask) | 0.87 | 0.88 |
| CC (volume) | 0.85 | 0.86 |
| CC (peaks) | 0.77 | 0.82 |
| CC (box) | 0.82 | 0.83 |
| **Model composition** |  |  |
| Non-hydrogen atoms | 16524 | 17223 |
| Protein residues | 2076 | 2082 |
| Nucleic acid | 12 | 18 |
| **Mean B factors (Å<sup>2</sup>)** |  |  |
| Protein | 53.45 | 33.82 |
| Nucleic acid | 74.64 | 49.73 |
| **R.m.s. deviations** |  |  |
| Bond lengths (Å) | 0.005 | 0.007 |
| Bond angles (°) | 0.574 | 0.629 |
| **Validation** |  |  |
| Molprobity score | 1.66 | 1.48 |
| Clashscore | 3.98 | 3.28 |
| Poor rotamers (%) | 2.93 | 2.10 |
| **Ramachandran plot** |  |  |
| Favored (%) | 97.38 | 97.39 |
| Allowed (%) | 2.62 | 2.46 |
| Disallowed (%) | 0 | 0.14 |

### FRET endoribonuclease assay

Nsp15 cleavage was monitored in real-time as described previously (19,20). Briefly, 6-mer substrates were labeled with 5’-fIuorescein (FI) and 3’-TAMRA, where TAMRA quenches FI and cleavage is measured by increasing FI fluorescence (5’-FI-AAxxxA-TAMRA-3’; x nucleotides varied among substrates) (see Supplementary Table 1). The substrate (0.8 μM) was incubated with Nsp15 (2.5 nM) in RNA cleavage buffer (20 mM Hepes pH 7.5, 75 mM NaCl, 5 mM MnCl_2_, 5 mM DTT) at 25°C for 60 minutes. Fluorescence was measured every 2.5 minutes using a POLARstar Omega plate reader (BMG Labtech) set to excitation and emission wavelengths of 485 ± 12 nm and 520 nm, respectively. Three technical replicates were performed for each condition, and the assay was repeated with at least two independent protein preparations. Prism (Graphpad) was used to calculate significant differences using Dunnett’s T3 multiple corrections test.

### Urea-PAGE endoribonuclease assay

Double fluorescently-labeled RNA substrates (5’-FI and 3’-Cy5, 500 nM) were incubated with Nsp15 (50 nM) in RNA cleavage buffer (20 mM Hepes pH 7.5, 150 mM NaCl, 5 mM MnCl_2_, 5 mM DTT, 1 u/μL RNasin ribonuclease inhibitor) at room temperature for 30 minutes, with samples collected at 0, 1, 5, 10, and 30 minutes. The reaction was quenched with 2x urea loading buffer (8M urea, 20 mM Tris pH 8.0, 1 mM EDTA). Due to the expected size of cleavage products and the size of bromophenol blue, loading buffer without dye was used. To monitor the gel front, a control lane of protein only with bromophenol blue was run. To generate a ladder, alkaline hydrolysis of the RNA was carried out for 15 min at 90°C using 1 μM RNA in alkaline hydrolysis buffer (50 mM sodium carbonate pH 9.2, 1 mM EDTA) and quenched with 2x urea loading buffer. The cleavage reactions were separated using 15%-20% TBE-urea PAGE gels and visualized with a Typhoon RGB imager (Amersham) using Cy2 (λ_ex_=488 nm, λ_em_=515-535 nm) and Cy5 (λ_ex_=635 nm, λ_em_=655-685 nm) channels.

### Mass spectrometry of RNA cleavage products

Mass spectrometry was performed as previously described (20). Briefly, the FRET RNA substrate of interest (0.8 μM) was incubated +/- Nsp15 (2.5 nM) in RNA cleavage buffer for 30 minutes at RT. For mass spectrometry analysis, the reaction was chromatographically separated with a gradient of buffer A (400 mM hexafluoro-2-propanol, 3 mM triethylamine, pH 7.0) and buffer B (methanol). Parallel reaction monitoring (PRM) analyses were included in the MS analyses with included masses of m/z 914.14; 923.14; 1463.42.

### Molecular dynamics simulations

Based on the RNA bound cryo-EM hexamer structure of Nsp15, the initial structure of Nsp15-AUA hexamer complex was prepared by manually introducing an adenine nucleotide at the B_-2_ position. Except for H250, all histidine residues were selected to be Nε protonated. Since the ring nitrogen atoms on H250 were found to make two strong hydrogen bonds with the phosphate backbone and the carbonyl oxygen of S294, H250 was assigned the positively charged doubly-protonated form. After introducing all protons using the TLeap module of Amber.18 (32), the Nsp15-AUA hexamer system was solvated in 68,849 water molecules, while 203 sodium ions and 125 chloride ions provided the 100 mM salt concentration and the charge neutralization. A separate Nsp15-AUA monomer system, isolated from the hexamer, was also subjected to molecular dynamics. The monomer assembly was solvated with 24,545 water molecules. There were 57 sodium ions and 44 chloride ions also in the monomer system. The hexamer system consisted of 240,229 atoms while the monomer system had 79,295 atoms. The boundaries of the water boxes were at least 15 Å away from any protein or RNA atoms.

After proper equilibration of each system over 30 ns under various conditions, the CUDA implementation of the PMEMD module of Amber.18 with the amino acid represented by the FF14SB force field was used to simulate unconstrained dynamics for 500 ns for hexamer and monomer systems at 2 fs time step and 300 K under constant pressure. The Amber FF14 RNA force field was used for the ribonucleotide trimer. The particle mesh Ewald method was used in dealing with long range Coulomb and van der Waals interactions. For each system, two additional 500 ns simulations were performed. The starting structures of the additional runs were selected from the 30 and 40 ns conformations of the primary simulation with the randomized initial velocities to simulate alternate trajectories. The MMGBSA module of Amber.18 was implemented in free energy estimations with the selection of 0.15M salt concentration and the default parameters (IGB = 5) in the Amber module. Since the trinucleotide was not bound to the binding site residues during the entire half a microsecond production runs in most systems, the energy calculations were performed for each 50 ns segments (with 50 samples selected at each nanosecond) separately for each trajectory. When calculating the residues interaction energies, only the values from 50 ns segments with bound trinucleotides were selected.

## RESULTS

### Cryo-EM reconstructions of Nsp15 bound to RNA in pre- and post-cleavage states reveal substrate binding interactions

To gain insight into RNA cleavage by Nsp15, we determined cryo-EM structures of Nsp15 with RNA bound in pre- and post-cleavage states (Table 1, Figure 1, and Figure 2). Given the similar active site arrangement and chemistry between Nsp15 and RNase A, we hypothesized that analogous to RNase A, there may be additional base specific binding pockets in Nsp15 (23,33). RNase A has multiple phosphate and base binding pockets that mediate the position of the base 5’ (herein referred to as B_-1_) and 3’ (herein referred to as B_+1_) to the scissile phosphate. Similar to Nsp15, the base binding pocket at the B_-1_ position in RNase A confers specificity to pyrimidines, through a well conserved threonine (S294 in SARS-CoV-2 Nsp15). RNase A prefers purines following the pyrimidine and this preference is mediated by interactions in the B_-1_ binding pocket.

**Figure 1.**
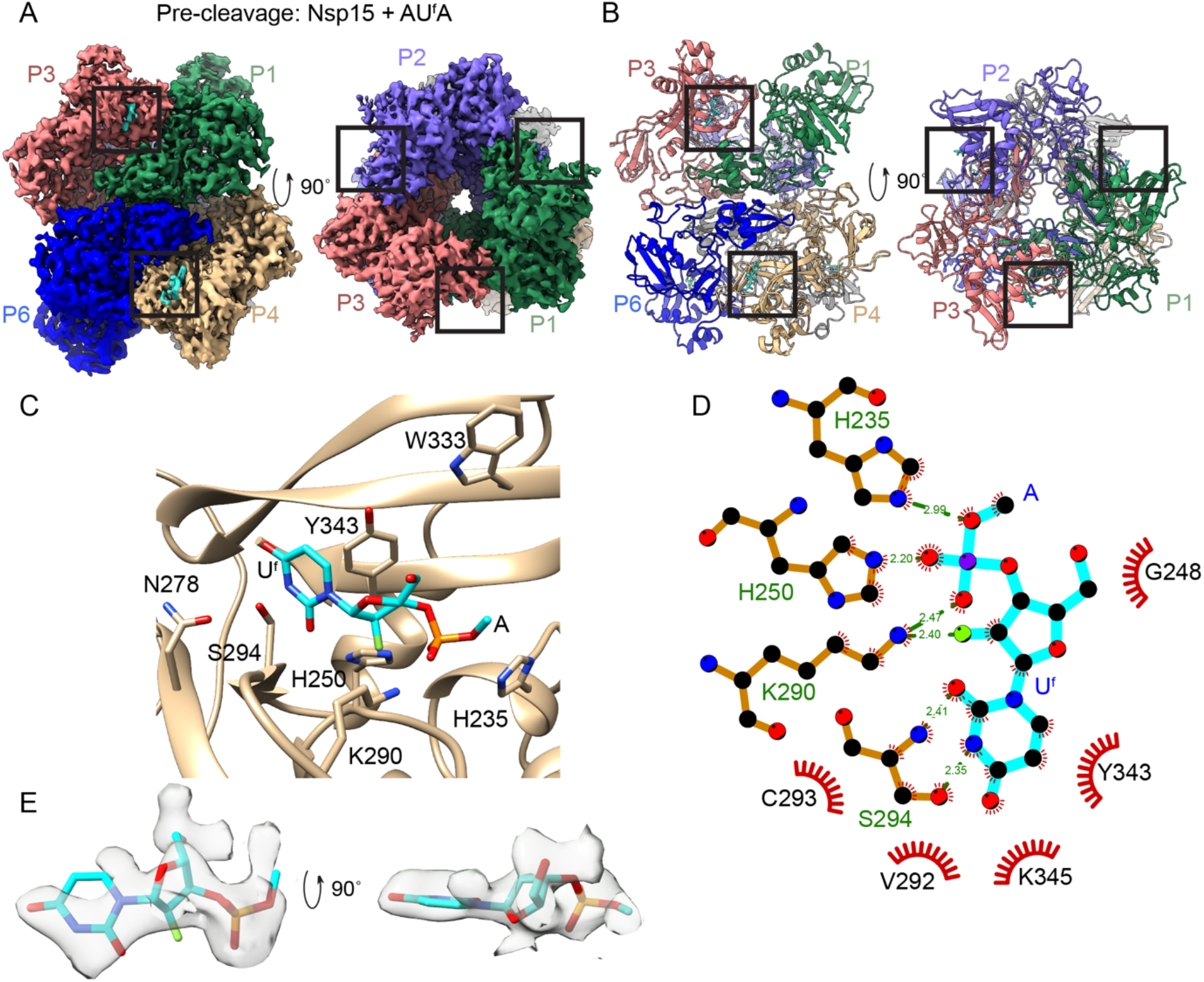
Pre-cleavage cryo-EM structure (Nsp15 + AU^f^A) shows the complete, uncut scissile bond. Side and top views of the cryo-EM map (**A**) and model (**B**). Protomers are colored and labeled. The active site is boxed in black and the RNA is colored teal. (**C**) Zoomed in active site with RNA bound. (**D**) 2D Ligplot (58) representation of interactions between the RNA and Nsp15. (**E**) The density for the AU^f^A ligand is shown.

To determine if Nsp15 has a B_-1_ binding pocket, we trapped Nsp15 in a pre-cleavage state by incubating Nsp15 with a modified RNA containing a 2’ fluorine substituting for the 2’-OH on the uridine ribose that prevents catalysis (AU^f^A; Supplementary Table 1) prior to vitrification. 2’ fluorine modifications have been well-characterized and exploited for generating ribonuclease resistant oligos (34,35); the small size of fluorine in particular makes it an ideal modification unlikely to disturb the structure. Cryo-EM data were collected using a Titan Krios microscope (Supplementary Figure 1 and Table 1). The pre-cleavage state map was determined to 2.5 Å resolution with C3 symmetry applied. RNA density within the EndoU active site was observed in three of the six active sites, corresponding to one of the two trimers in the C1 map; poor resolution of the EndoU domains of the other trimer hindered conclusive identification of RNA within that trimer. We did not observe any asymmetric features within the trimers so C3 symmetry was subsequently applied due to the resolution enhancement from effectively tripling the particle number. Overall, the structure of the pre-cleavage state is very similar to previously determined SARS-CoV-2 Nsp15 structures bound to mono or dinucleotides, with RMSD values of <1 Å (Figure 1 and Supplementary Figure 1). We were able to visualize the complete, uncut scissile phosphate of the tri-nucleotide substrate (AU^f^A), although only the 2’-fIuoro-uridine and the ribose C5’ of the B_+1_ A could be modeled in the density (Figure 1C and 1E). The fluorine was properly oriented within the catalytic triad (H235/H250/K290) even though no chemistry could take place; this also allowed for the U 3’-PO_4_ to be positioned for hydrogen bonding with the catalytic triad. Consistent with UMP- and UTP-bound structures, the uracil base is poised to form hydrogen bonds with S294, such that Nsp15 can discriminate between U and other bases, while the uridine ribose group is oriented with Y343 to form van der Waals interactions (Figure 1D and Supplementary Figure 2) (20,21). Thus, this positioning represents a uridine poised for cleavage. The lack of well resolved density for either adenine base suggests that in contrast to RNase A, beyond the uridine recognition site Nsp15 does not have strong secondary base binding sites.

To further probe RNA recognition by Nsp15 we determined the cryo-EM structure of Nsp15 in the post-cleavage state. We captured the post-cleavage state by incubating Nsp15 with excess unmodified RNA (AUA) prior to vitrification and cryo-EM data collection. Use of an unmodified RNA had the potential to lead to the capture of multiple states of Nsp15 (pre-cleavage, cyclic phosphate intermediate, and mono-phosphate product), however during classification and refinement we were only able to identify a single state containing the 5’ final product (AU-3’P). Inspection of the C1 map revealed unambiguous density for RNA in all 6 active sites, however the RNA density was better resolved in one of the two trimers, so C3 symmetry was applied. The final cryo-EM reconstruction of the post-cleavage state went to 2.2 Å resolution and the map contains well resolved side chain density throughout the entire molecule. Analogous to the pre-cleavage state the overall structure of the post-cleavage state is similar to previous Nsp15 structures, with RMSD values of <1 Å (Figure 2 and Supplementary Figure 3), suggesting that there are no large conformational changes following transesterification and hydrolysis.

**Figure 2.**
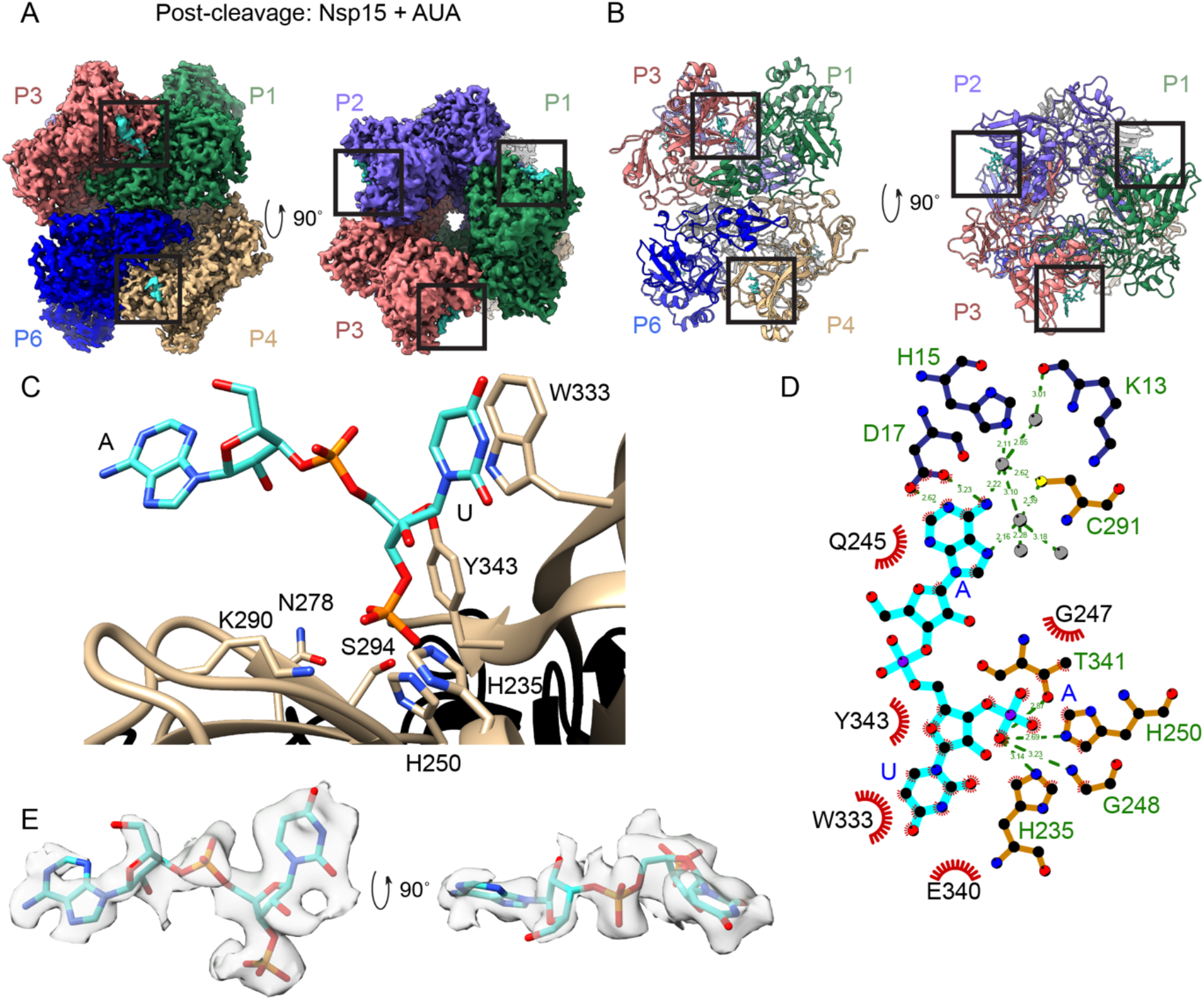
Post-cleavage cryo-EM structure (Nsp15 + AUA) reveals a change in U positioning. Side and top views of the cryo-EM map (**A**) and model (**B**). Protomers are colored and labeled. The active site is boxed in black and the RNA is colored teal. (**C**) Zoomed in active site with RNA bound. (**D**) 2D Ligplot (58) representation of interactions between the RNA and Nsp15. (**E**) The density for the AUA ligand is shown.

The RNA density within the active site was modeled as AU-3’P RNA, which represents the final 5’ product following transesterification and hydrolysis (Figure 2C and 2E). The high-resolution density for the uracil base is unambiguous. Notably, the position of the uracil base and ribose differs from the pre-cleavage state and the majority of previously determined nucleotide or nucleotide analogue structures of Nsp15. While the 3’-PO_4_ remains in the same place as the pre-cleavage state, the uracil has moved to pi-stack with W333 instead of interacting with Y343 and S294, a >10 Å movement of the base (Figure 2D). The observed movement of the uracil in the post-cleavage state aligns with both a recently published crystal structure of Nsp15 bound to 3’ UMP (PDB ID 6X41, (21)) and a deposited crystal structure of Nsp15 bound to uridine 3’,5’-diphosphate (PDB ID 7K1O), which represent a minimal product nucleotide (Supplementary Figure 2D). Those structures, along with our postcleavage state, are the only Nsp15-substrate structures where the nucleotide substrate contains a free 3’ PO_4_. The alignment seen between these structures provides further support that the repositioning of the uracil base following cleavage is a relevant post-cleavage state, though it may be one of several states sampled post-cleavage (Figure 3). While it is unclear why the uracil would leave the uracil binding pocket this structure suggests that W333 could be another critical residue for RNA binding within Nsp15. The role of W333 in RNA binding is further supported by a recent crystal structure of Nsp15 bound to a 5’ GpU dinucleotide in which the guanine base pi-stacks with W333 (21). We also noticed the position of S294 flips from the position it uses to form a hydrogen bond with the uracil base when it sits in that pocket. The result of this movement is that S294 moves out of hydrogen bonding distance from N278.

**Figure 3.**
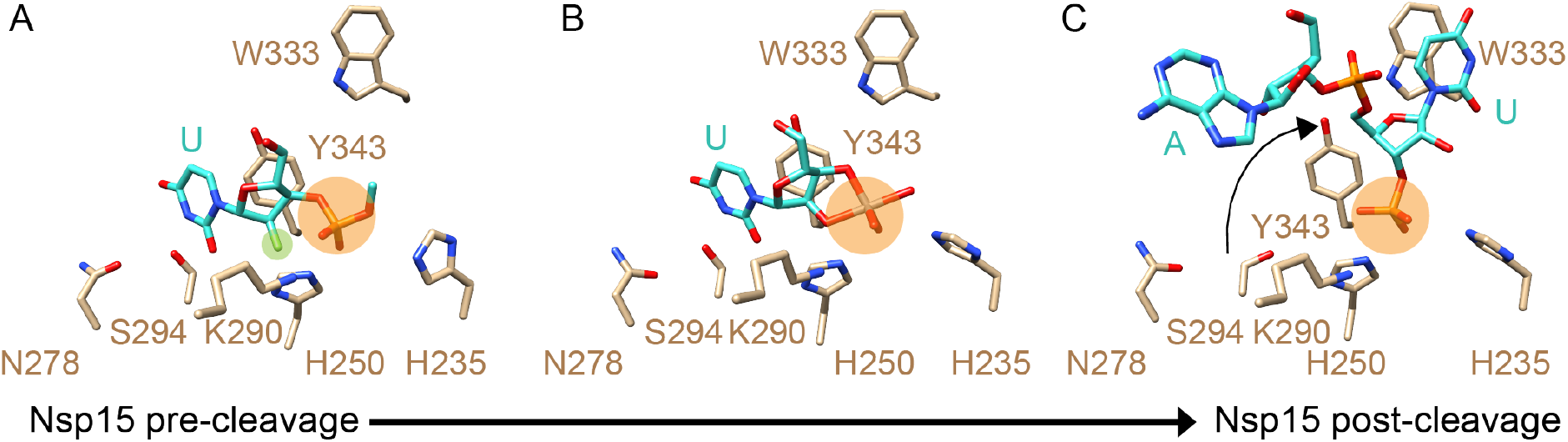
Nsp15 structures across the complete reaction mechanism. In all panels, active site residues of interest are shown as tan sticks, the RNA as turquoise sticks, and the scissile phosphate highlighted in orange. (**A**) The pre-cleavage structure with U^f^* (highlighted in green) poised for cleavage (this work). (**B**) The cyclic intermediate state determined using the mimic uridine-2’,3’-vanadate (PDB: 7K1l (21)). (**C**) The post-cleavage structure with the U pivoting to pi-stack with W333 (this work).

In contrast to the pre-cleavage state, we also observed weak density corresponding to the adenine base from the AU-3’P product. The density for the adenine is less well defined than the uridine but lies adjacent to the N-terminal domain (NTD) from a neighboring Nsp15 protomer (Figure 2C and 2D). The majority of these interactions are mediated through water molecules, although D17 does make base specific interactions with the adenine. Intriguingly, RNA crosslinking and mutational analysis with SARS-CoV-1, identified this same region of the NTD as being important for RNA binding (36). While the adenine density is poor the Nsp15 side chain density was well-resolved, and we observed unexpected density near H15 and C291 in close proximity to the active site (Supplementary Figure 5A and 5B). The density is continuous with the cysteine side chain suggesting a putative cysteine modification. Analysis of our recombinant Nsp15 by mass spectrometry suggests this could be a βME adduct, although a full adduct does not fit into the observed density (data not shown). However, we cannot rule out the possibility that the βME modification occurred during sample preparation for mass spectrometry as mass spectrometry also identified a βME modification on C293, an interior facing residue with no extra density in our map. Due to this ambiguity, the additional density was modeled as water molecules but this observation suggests that C291 could be a reactive cysteine. Collectively the structures of Nsp15 in pre- and post-cleavage states revealed new insight into RNA recognition that prompted us to examine the function of residues surrounding the active site.

### Molecular dynamics simulations and energy calculations support cryo-EM structural observations

Our cryo-EM structure provided partial density for the uncleaved trinucleotide RNA substrate, with the nucleotides adjacent to the uridine seemingly highly dynamic. Therefore, we turned to molecular dynamics simulations to further characterize the behavior of the nucleotides near the active site. Root mean square deviations (RMSDs) were used to establish the stability of the simulated systems (Supplementary Figure 4A) in which the isolated monomer systems displayed elevated dynamics (as assessed by RMSDs) compared to the protomers assembled into the hexamer. This is consistent with our previous molecular dynamics simulations revealing that the hexamer is important for protein stability (20). To assess RNA trinucleotide binding to Nsp15, we monitored multiple distances during the dynamic simulations, including three distances from the catalytic triad (K290, H235, H250) and two distances from the uridine discriminator (S294): NZ(K290)-P(PO_4_); NE2(H235)-O5’(B_+1_); NE2(H250)-O(PO_4_); N3(S294)-O2(U); OG(S294)-N3(U). These distances (Supplementary Figure 4B) guided us in deciding whether or not the AUA trinucleotide was bound, leading to subsequent analyses of dynamics behavior and related interaction strengths at the residue level. The thermal fluctuations were evaluated through B-factors calculated from the simulations (Figure 4); much larger fluctuations were displayed for the bases of both neighboring nucleotides compared to the uridine. However, assembly into a hexamer reduced the range of B-factors (Figure 4, bottom panel). The monomer dynamics show the possible independent movements among NTD, Middle, and EndoU domains in both B-factors and the dynamic cross correlations (Supplementary Figure 4C).

**Figure 4.**
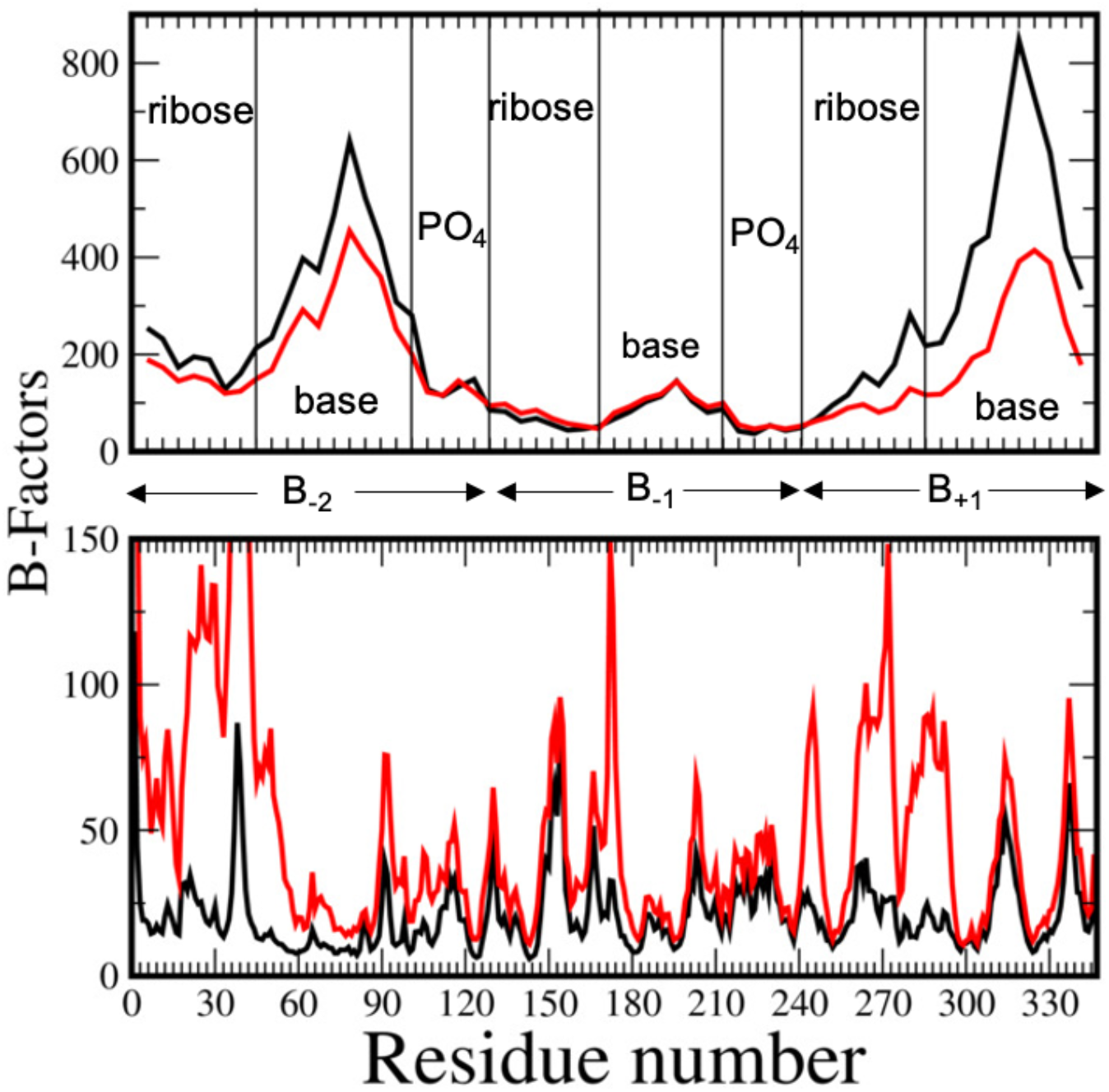
Thermal fluctuations (B-factors) calculated from the molecular dynamics simulations show increased movement for B_-2_ and B_+1_ compared to the B_-1_ position. Conformations in which the central U was found to be in a bound conformation were selected for the B-factor calculations from all three simulations for each system. Atomic B-factors of the RNA trinucleotide are shown in the top panel (hexamer – black; monomer – red). The backbone B-factors calculated by averaging the atomic B-factors of backbone heavy atoms of each residue for hexameric (black) and monomeric (red) systems are shown in the bottom panel. Each monomer in the hexamer was used to evaluate these averages from all three runs.

We then used MM/GBSA (molecular mechanics/generalized Born surface area) analysis to evaluate the contributions of Nsp15 active site residues to binding free energies. Residues providing binding energies with magnitudes above a 0.5 kcal/mol threshold are listed in Supplementary Tables 2 and 3. K290 and H250 provide the biggest contribution for the binding of the RNA trinucleotide, further highlighting the importance of the B_-1_ phosphate for driving nucleotide binding. Similar characteristics were observed in the structurally similar RNase A system (22). Aside from the phosphate in the active site, the residues with the strongest energetic contributions mainly do so through interactions with the base and ribose of the U. Residues interacting with the B_+1_ base overall feature larger energy values than the B_-2_ base (with the exception of W333), suggesting that while both bases are not as fixed as U, the B_+1_ base has more favorable interactions with Nsp15 active site residues (Supplementary Tables 2 and 3).

### Nsp15 active site mutants exhibit reduced or abrogated RNA cleavage

To determine the significance of SARS-CoV-2 Nsp15 active residues in mediating RNA interactions and supporting cleavage we made a series of single point mutations. We expressed and purified the following Nsp15 active site variants: N278A, S294A, W333A, and Y343A, which are largely well-conserved across coronaviruses (Figure 5A and Supplementary Figure 6). All four active site variants were purified as stable hexamers, indicating that they did not disrupt the oligomerization of Nsp15 (Supplementary Figure 5). We measured RNA cleavage with a FRET-based assay (19,20) using 6-mer RNA substrates with 5’ fluorescein and 3’ TAMRA labels. TAMRA quenches the fluorescein signal of the uncleaved RNA, therefore increasing fluorescence over time serves as a readout for cleavage. We used three different RNA substrates for this assay to address Nsp15’s ability to cleave RNA following either a U or a C, with a 6-mer polyA RNA serving as the negative control. The S294A and Y343A active site variants lack appreciable nuclease activity (Figure 5C). This confirms the critical importance of Y343 in orienting the uridine ribose and S294 in hydrogen bonding with the uracil base. S294 and Y343 are conserved in SARS-CoV-1 and the results of our mutational analysis are in good agreement with earlier studies with SARS-CoV-1 Nsp15 (36). Mutating N278A, the residue that interacts with S294 to position it correctly, resulted in hexameric enzyme that was ~2-fold less active and did not discriminate between U and C as strongly as WT (Figure 5C). At the endpoint of this assay, WT Nsp15 cleaved 40x more U/C, while Nsp15 N278A only cleaved 3x more U/C. Thus, N278 is important for maintaining uridine preference in SARS-CoV-2 Nsp15. Similar to N278A, the W333A variant was ~2-fold less active than WT Nsp15. This suggests that while W333 can provide a platform for pi-stacking with RNA bases, this interaction is not absolutely critical for RNA cleavage.

**Figure 5.**
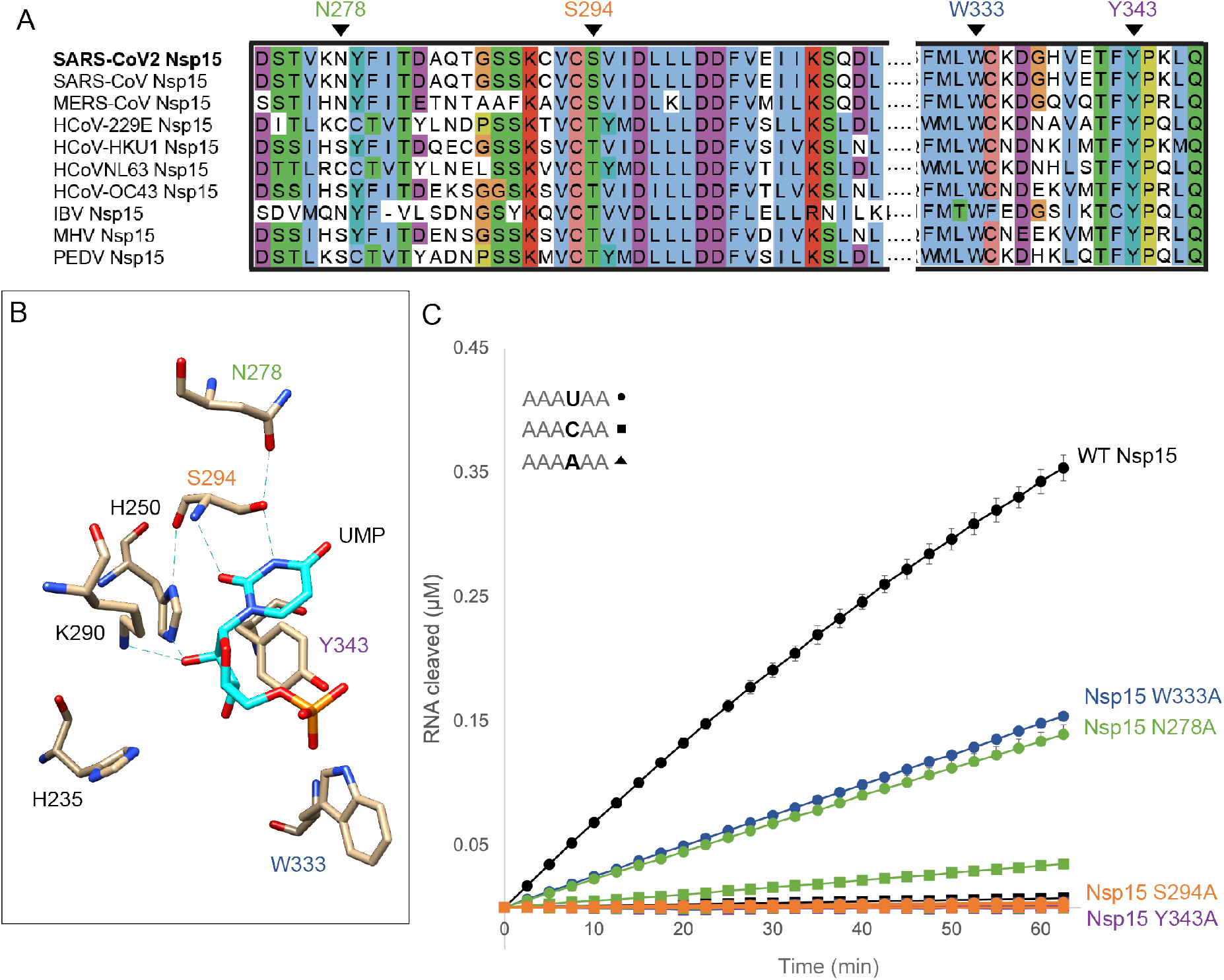
Nsp15 active site mutants reveal roles of critical residues. (**A**) Sequence alignment for selected regions of the Nsp15 EndoU domain across human coronaviruses and several major animal coronaviruses. Mutated residues are labeled. Residues that were mutated are colored to match the corresponding reaction curves. (**B**) Active site model of a pre-cleavage state (mimicked with 5’ UMP, PDB: 6WLC) depicting the catalytic triad and other important active site residues. (**C**) FRET time course data for Nsp15 active site mutants. Nsp15 WT and mutants (2.5 nM) were incubated with RNA (0.8 μM) at room temperature and fluorescence was monitored every 2.5 minutes for an hour. The average of a representative technical triplicate is plotted with standard deviation error bars. At least two biological replicates were performed for each mutant. Three different substrates were tested for each mutant: AAA**U**AA (circles), AAA**A**AA (squares), and AAA**C**AA (triangles). Each mutant is represented by a different color: WT Nsp15 (black), Nsp15 W333A (blue), Nsp15 N278A (green), Nsp15 S294A (orange), and Nsp15 Y343A (purple).

We also determined the importance of several residues near the weak adenine (B_-2_) base interaction site observed in the post-cleavage state, which lies at an interface with the EndoU domain of one protomer and the NTD of another protomer within the same trimer. Given that we noticed extra density extending from the C291 side chain, we tested the importance of this residue in mediating cleavage (Supplementary Figure 5). Mutating C291A did not significantly affect the oligomerization or activity of Nsp15, which is not surprising given that it is not well conserved (only SARS-CoV-1 also has a cysteine at that position; Supplementary Figure 6). However, mutating the nearby H15A (from a neighboring protomer) disrupted the oligomerization of Nsp15 and resulted in predominantly inactive, monomeric enzyme (Supplementary Figure 5D, 5E and 5F). Previous work with SARS-CoV-1 Nsp15 revealed that the first 25 residues of the NTD were important for oligomerization, as a truncation produced monomeric Nsp15 (36,37). Thus, mutation of H15 which lies in a hinge region prior to a helix that interacts with the neighboring protomer, may be disrupting the folding or packing of the NTD.

We mutated two additional residues near the B_-2_ base from the NTD including K13A and D17S, (Fig 2D) and saw no significant change in oligomerization compared to WT Nsp15 (Supplementary Figure 5D and 5E). However, activity of K13A was ~2-fold lower, while activity of D17S was ~2-fold higher suggesting the neighboring NTD does affect EndoU function (Supplementary Figure 5F). One possible explanation for these results is charge stabilization of the RNA, since overall the EndoU surface is very negative. The robust extra density near C291 and H15 includes a shorter distance between atoms reminiscent of a metal ion interaction. Manganese ions are important for Nsp15 activity, but have never been localized in the active site, suggesting they are not structural metals. One hypothesis is that Mn^2+^ ions provide charge shielding to increase the favorability of the RNA-Nsp15 interaction, and thus are more transient. Given that loss of the K13 positive charge decreased activity, and the loss of the D17 negative charge increased activity, our data may point to a role in charge stabilization for this region of the active site. Overall, the identification of these new active site residues from a neighboring protomer important for Nsp15 activity provide more evidence for allosteric regulation and suggest a new way to inhibit this enzyme by targeting the NTD/EndoU interface. All together our structural and functional analysis highlights the significance of several Nsp15 residues in mediating RNA cleavage, specificity, and oligomerization.

### *In vitro* FRET endoribonuclease assay reveals Nsp15 prefers a purine in the position following the uridine

While the structures of Nsp15 in the pre-and post-cleavage states did not reveal strong secondary base binding sites, RNA cleavage assays revealed that Nsp15 has a preference for a purine 3’ to the uridine in the cleavage site. Using the FRET endoribonuclease assay described above, we studied the importance of the bases in the −2 and +1 position (B_-2_, B_+1_) relative to the uridine being cleaved (B_-1_) using a series of oligomers with a single nucleotide change (Supplementary Table 1 and Figure 6A). Substituting an A or C for U in the B_-1_ position resulted in minimal cleavage, confirming Nsp15’s specificity for cleavage following a U. In contrast, substitutions in the B_-2_ position did not significantly affect Nsp15 cleavage. However, substituting a C in the B_+1_ position resulted in significantly less cleavage compared to an A or G (Figure 6B). We further looked at the effect of substituting a U in either the B_-2_ or B_+1_ position. Because the presence of two U’s introduces a second cleavage site, we measured time courses with either U or a modified uncleavable U in the B_-2_ or B_+1_ position. Addition of a U in the B_-2_ position does not impact cleavage significantly but addition of a U in the B_+1_ position led to a reduction in cleavage, albeit not as significant of a reduction as with a C (Supplementary Figure 7A). We further analyzed the cleavage products of the unmodified double U RNAs by mass spectrometry which showed the accumulation of cleavage products following the uridine nucleotides at all three positions (Supplementary Figure 7B and 7C). Therefore, the differences observed in cleavage with the double U constructs suggests that the position of the U relative to the 5’ or 3’ end may also impact cleavage. Based on these endoribonuclease assay results, we define the consensus motif for Nsp15 cleavage as (N)(U)^(R>U≫C) (where N is any nucleotide and R is a purine). Together these results reveal that Nsp15 has a preference for purines over pyrimidines in the B_+1_ position, however Nsp15 can efficiently process substrates with tandem U’s.

**Figure 6.**
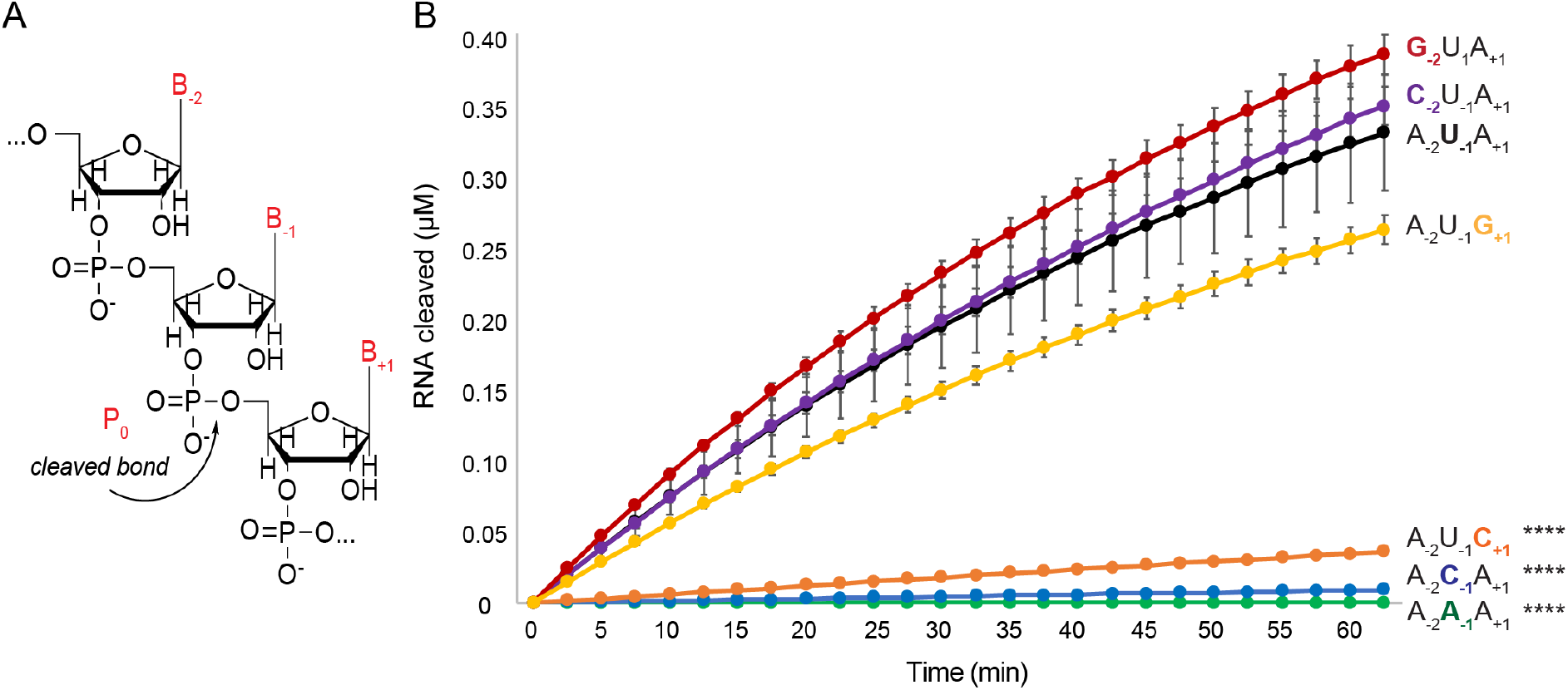
Nsp15 RNA specificity identified by a FRET endoribonuclease assay reveals the importance of the B_+1_ position. (**A**) Illustrative RNA diagram highlighting the scissile phosphate bond (Po), and defining the nomenclature of the bases of interest (B_-2_, B_-1_, B_+1_). (**B**) Reaction curves for comparing cleavage of different RNA sequences. WT Nsp15 (2.5 nM) was incubated with RNA (0.8 μM) at room temperature and fluorescence was monitored every 2.5 minutes for an hour. The average of a representative technical triplicate is plotted with standard deviation error bars. At least two biological replicates were performed for each mutant. For simplicity, only the three bases that were changed in this experiment are listed by the respective curves. U in the B_-1_ position with A in the B_-2_ and B_+1_, is shown in black. B_-1_ was changed to an A (green) and C (blue). B_-2_ was changed to G (red) and C (purple). B_+1_ was changed to G (yellow) and C (orange). Dunnett’s T3 multiple corrections test was performed using Prism for pairwise comparisons. The p<0.0001 for the indicated (****) curves.

### Cleavage of biologically relevant substrates shows Nsp15 has a clear preference for U^A versus U^C

Our FRET-based cleavage assays revealed that Nsp15 demonstrates selectivity beyond uridine recognition in small 6-mer substrates leading us to ask whether this specificity is conserved in a) longer and b) more biologically relevant RNA substrates. Cyclic phosphate sequencing of MHV (a mouse model coronavirus) infected macrophages revealed MHV Nsp15 cleavage products throughout the positive strand of the MHV genome, including in conserved transcriptional regulatory sequences (TRS) (13). TRS sequences are short, conserved motifs that lead to discontinuous viral RNA synthesis to produce sub-genomic RNAs (38). The consensus TRS body (TRS-B) sequence from SARS-CoV-2 precedes each gene and promotes template switching by hybridizing to the TRS leader (TRS-L). The core TRS-B sequence includes one uridine and several cytidines, making it a putative substrate for Nsp15. We synthesized ~20-mer oligos containing the consensus TRS-B and flanking sequence for the nucleoprotein (N) as well as the spike (S) protein sub-genomic RNAs with labels on both the 5’ and 3’ ends (Table S1). We analyzed these oligos with the RNAfold server to confirm that the short constructs used in our assays are unlikely to form secondary structures (39).

We incubated WT Nsp15 with the TRS-N and TRS-S containing RNA substrates and then resolved the cleavage products on denaturing urea gels. We observed over time the accumulation of cleavage products following the uridines in both the TRS-N and TRS-S substrates (Figure 7). The time courses are consistent with the FRET-based cleavage data, although we had to use a higher ratio of protein to RNA to be able to visualize the products well. These higher protein concentrations are in-line with other Nsp15 activity assays (40). This data demonstrates that WT Nsp15 prefers purines in the B_+1_ position in longer substrates as we observed for the six nucleotide substrates. There are four U’s within the TRS-N sequence with differing B_+1_ nucleotides: U_1_^C, U_4_^C, U_6_^A and U_19_^A. We observed faster accumulation of the U_6_^A cleavage product over the U_1_^C and U_4_^C products. Cleavage at U_19_^A was minimal, an effect that could be due to a bias for shorter products in this assay and/or issues detecting smaller Cy5-products (41,42). The TRS-S sequence contains a single U_7_^A cleavage site. In addition to the U7 cleavage product observed with the TRS-S substrate, we also detect a small amount of cleavage following C_3_^A and C_6_^U. We repeated these cleavage assays with the Nsp15 N278A variant that demonstrated change in U/C specificity in our FRET cleavage assay (Fig. 4). Consistent with our assay results with a six nucleotide substrate, the N278A Nsp15 variant cleaved more slowly and produced more C cleavage products than WT Nsp15 (Fig. 7). We further tested the B_+1_ purine preference using another 20-mer oligomer (Table S1) that was previously used to measure cleavage in SARS-CoV-2 Nsp15 (21) featuring multiple uridines with different B_+1_ bases. The time course again shows that the products accumulate preferentially, with the product resulting from a purine in the B_+1_ position (U_8_^G) predominating (Supplementary Figure 8). Collectively the results from our gel-based cleavage assays support the results from our FRET cleavage experiments that Nsp15 prefers to cleave U’s following purines.

**Figure 7.**
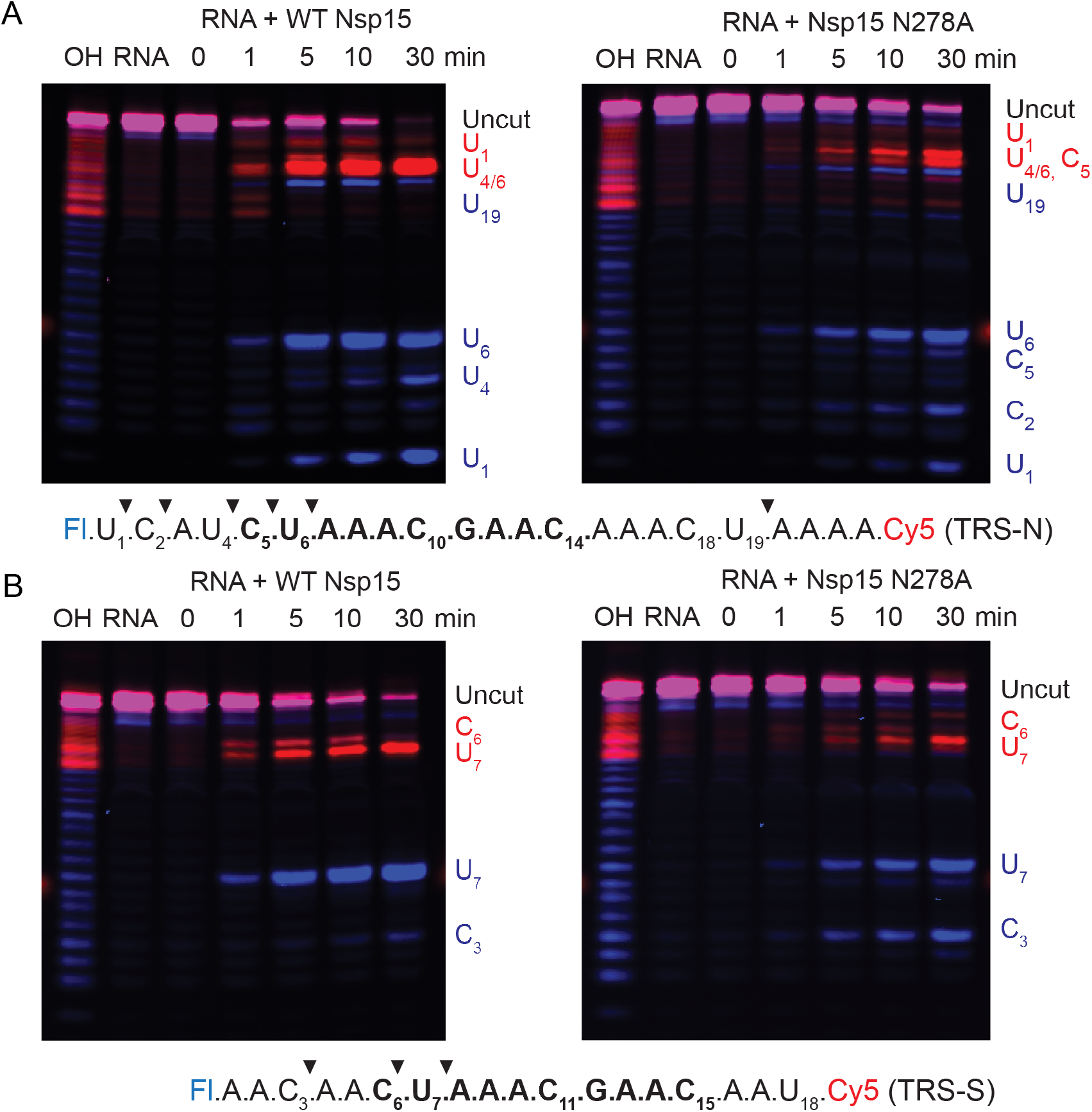
Cleavage gels of biologically relevant TRS sequences show B_+1_ sequence preference. The sequence is shown below the gels with the conserved consensus sequence in bold and the labels colored to match the overlays. Nsp15 (50 nM) was incubated with 5’-FI-RNA-Cy5-3’ (500 nM) for 30 minutes at room temperature. Time course samples were taken at the indicated times and quenched with loading buffer. Alkaline hydrolysis of the RNA was performed to produce a ladder (lane OH) Representative gels are shown from three technical replicates. (**A**) Cleavage gels for WT Nsp15 (left) and Nsp15 N278A (right) with TRS-N. Images for the 5’ (blue) and 3’ (red) products were overlaid. (**B**) Cleavage gels for WT Nsp15 (left) and Nsp15 N278A (right) with TRS-S. Images for the 5’ (blue) and 3’ (red) products were overlaid.

Finally, we also looked at Nsp15’s ability to degrade polyU sequences and found that Nsp15 efficiently degrades polyU containing RNAs in vitro. In addition to having numerous targets across the positive strand, previous work with MHV Nsp15 demonstrated activity in the PUN tract of the negative strand RNA *in vitro* and *in vivo* (9). We assessed the ability of SARS-CoV-2 Nsp15 to cleave polyU sequences in two ways. First, we performed a cleavage assay with a 5’ fluorescein labeled U17 oligomer. Nsp15 acted efficiently on this substrate, leading to an accumulation of the single nucleotide product over time (Supplementary Figure 9). Similar to a previous study with MHV-CoV-1 Nsp15 (9), we also tested substrates with the SARS-CoV-2 wild-type negative strand sequence with and without the 5’-polyU sequence, as well as oligomers with the negative strand changed to have no uridines or no pyrimidines, with 5’-fluorescein and 3’-Cy5 labels (Figure 8, Supplementary Figure 10 and Supplementary Table 1). The dual labels allowed us to track formation of both 5’-polyU products and the 3’-negative strand products. Similar to the polyU alone cleavage, Nsp15 efficiently degraded the entire PUN RNA sequence at the 5’ end. In addition to cleaving the PUN, Nsp15 cleaved uridines within the negative strand. When we removed the U’s from the negative strand, a small amount of C cleavage products were observed, whereas removal of all pyrimidines resulted in an RNA that was not cleaved by Nsp15. These results are in excellent agreement with the earlier work showing that Nsp15 targets the PUN RNA sequence, however Nsp15 activity is not restricted to the PUN as it also cleaves additional uridines within the negative strand. Our structural, molecular dynamics, and biochemical data combined demonstrate that Nsp15 activity is driven by interactions with uridines in viral RNA. Thus, our current model features the six active sites cleaving U’s efficiently, in both the 5’ polyU sequence of the negative strand and throughout the rest of the genome. This occurs regardless of the surrounding sequence, although our data show Nsp15 does prefer a purine in the B_+1_ position outside of the negative strand polyU sequence.

**Figure 8.**
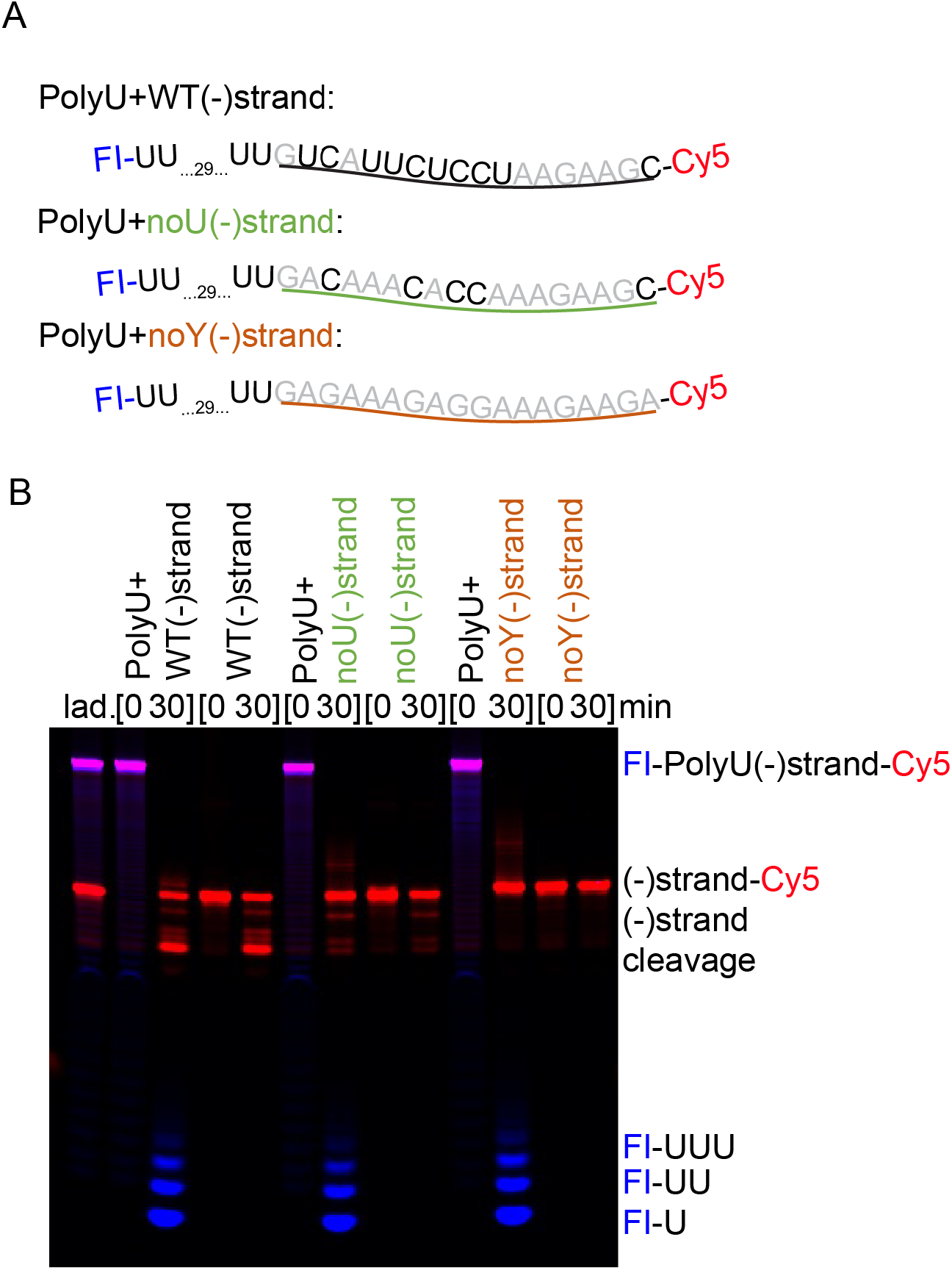
Cleavage gels of negative strand sequences with both the PUN and internal Us shows Nsp15 acts across the sequence. Nsp15 (50 nM) was incubated with RNA (500 nM) for 30 minutes at room temperature. The polyU-NS substrates had 5’-FI and 3’-Cy5 labels, while the NS substrates only had 3’-Cy5 labels. (**A**) Cartoon depiction of the substrates tested. (**B**) Summary cleavage gel for the time course reactions showing only the 0- and 30-minute samples for each substrate.

## DISCUSSION

Despite its critical role in coronavirus pathogenesis, how Nsp15 recognizes its RNA targets was poorly understood. Here we characterized the cleavage motif of SARS-CoV-2 Nsp15 to provide insight into how Nsp15 processes viral RNA. We determined cryo-EM structures of Nsp15 in the pre- and post-cleavage states. While we hypothesized that Nsp15 would contain additional base binding sites analogous to RNase A, we did not observe any strong secondary base binding sites in our cryoEM reconstructions, which suggests that beyond the uridine binding pocket the rest of the RNA is not fixed in place by strong interactions, and therefore averaged out during data processing. We did observe some density for the B_-2_ base preceding the uridine in the post-cleavage reconstruction towards the edge of the active site and adjacent to the NTD of a neighboring protomer. We identified several NTD residues from the adjacent protomer that interact with the B_-2_ adenine in our model and are important for oligomerization and nuclease activity. Therefore, in addition to being a more stable assembly (20,21), the hexamer may also be necessary for the NTD to support engagement of RNA in the active site. Thus, our structure-based point mutations suggest Nsp15 could be inhibited by disrupting the EndoU/NTD interface at the edge of the active site, which should destabilize the hexamer and lead to inactive monomeric enzyme.

While previous work established the role of S294 in conferring uridine specificity (20,21,36), here we identified N278 as a key residue in maintaining uridine preference in SARS-CoV-2 Nsp15. This residue is identical in SARS-CoV-1 and MERS Nsp15, but is only somewhat conserved (i.e. polar) across other coronaviruses that cause human disease (Supplementary Figure 6). Our structures show that N278 interacts with S294 to orient the hydrogen bond network to select uridines. The N278A variant showed reduced activity on a single U-containing substrate, but enhanced activity on a single C-containing substrate in comparison with WT Nsp15. The conservation of S294 shows a similar pattern to N278; the equivalent residue is a serine in SARS and MERS Nsp15, but is a threonine in the other human coronaviruses. Similarly, MHV Nsp15 has a serine at the equivalent position to N278, and a threonine equivalent to S294 (Supplementary Figure 6). Cyclic phosphate sequencing of MHV-infected cells revealed that MHV Nsp15 cleaves both U^A and C^A sequences (13), which differs from our in vitro activity assays with SARS-CoV-2 Nsp15 that reveal a clear preference for cleaving 3’ of U over C. While we cannot exclude the possibility that other viral or host factors may influence the sequence specificity of Nsp15 which could give rise to non-uridine cleavage products, our data suggests that the N278/S294 pair in SARS-CoV-2, SARS-CoV-1, and MERS may lead to a stronger preference for cleaving uridines.

We also probed substrate specificity via *in vitro* cleavage assays and found SARS-CoV-2 Nsp15 exhibited robust cleavage of PUN RNA, but appeared to prefer purines 3’ of the cleaved uridine in sequences outside of the polyU tract. Thus, Nsp15 appears to have evolved to cleave uridines rather indiscriminately. Interestingly, the base composition of coronaviruses is heavily biased towards uridines (43), which suggests both a regulatory interplay and an additional need to target Nsp15 so it is not acting in an uncontrolled fashion. For example, the nascent viral RNA destined to be packaged must be protected from Nsp15 activity. Additionally, Nsp15 activity towards TRS sequences suggests it could play a role in regulating transcription, since TRS sequences play an important role in coronavirus transcription.

There are several putative mechanisms for regulation of Nsp15 nuclease activity. Modification and/or secondary structure of the viral RNA could regulate nuclease activity. For example, coronaviruses encode a 2’-O-methyltransferase (Nsp16); the modification produced by this enzyme would be expected to hinder Nsp15 endoribonuclease activity (38,44). SARS-CoV-1 Nsp15 does not cleave 2’-methylated RNA substrate efficiently and preferentially cleaves unpaired U’s within a structured RNA substrate (36). Recent papers have mapped the secondary structure of the SARS-CoV-2 genome and found evidence for highly stable RNA folding throughout the genome (45–49). Beyond RNA modification and secondary structure, another potential mechanism of Nsp15 nuclease regulation is through compartmentalization. Extensive membrane rearrangement occurs within cells during coronavirus infection, leading to the formation of double membrane vesicles and other convoluted membrane structures (50–52). These membrane rearrangements are thought to help hide the transcription intermediate, double stranded RNA from host cell anti-viral sensors. This could lead to a local concentration of viral enzymes and viral RNA so that viral RNA editing is favored over host RNA editing. Recent work demonstrated Nsp15 endoribonuclease activity also hindered the formation of anti-viral stress granules by regulating the formation of viral dsRNA (14). There is also evidence that Nsp15 may inhibit autophagy (53). Thus, the cellular localization of Nsp15 could also affect its targeting. Finally, other viral proteins may influence Nsp15 RNA targets and regulation in host cells, as Nsp15 is believed to localize within the replication-transcription complex of Nsps, including the RdRp complex (54,55). Stable or transient interactions with other RTC proteins could thus direct Nsp15 specificity and activity.

Overall, this work establishes SARS-CoV-2 Nsp15 as a largely non-specific endoribonuclease with recognition for a minimal consensus motif (N)(U)^(R>U≫C) (where N is any base and R is a purine). Our data show that Nsp15 acts in a distributive fashion to catalyze cleavage following uridines. Nsp15 is a key player in blocking activation of host dsRNA sensors by preventing the accumulation of viral RNA and a promising therapeutic target (9,13,14). Prokaryotic and eukaryotic RNase A and EndoU family proteins have been characterized as distant homologues, with extensive bioinformatics revealing new members of the families throughout the kingdoms of life (56). These two families share a catalytic triad that represents an ancient RNA processing mechanism, which has been adapted to function in host defense and innate immunity in many instances (57). This work reveals that similar to RNase A, Nsp15 is a broad-spectrum endoribonuclease primarily guided to its cleavage targets by recognition of a single uridine. Our integrated structural and functional characterization of Nsp15 provides a platform for the development of innovative therapeutic strategies against coronaviruses.

## DATA AVAILABILITY

Atomic coordinates and structure factors for the reported cryo-EM structures have been deposited with the Protein Data Bank (PDB) and the Electron Microscopy Data Bank (EMDB) under accession numbers EMDB-24137, EMDB-24101, PDB ID: 7N33, and PDB ID: 7N06.

## SUPPLEMENTARY DATA

Supplementary Data are available with the manuscript.

## ACKNOWLEDGEMENT

We would like to thank Dr. Rick Huang and Allison Zeher for help with cryo-EM data collection. This work utilized the NCI/NICE Cryo-EM Facility. We would like to thank all members of the Stanley Lab and NIEHS cryo-EM core facility for helpful discussions on this project. Finally, we would like to thank Drs. Traci Hall and Oswaldo Lozoya for their critical reading of this manuscript.

