## Supplemental Information for "Characterization of SARS2 Nsp15 Nuclease Activity Reveals it’s Mad About U"

This PDF Includes:

Supplementary Tables: S1 to S2

Supplementary Figures: S1 to S10

Supplementary References

**Supplementary Table 1. List of RNA oligos used in this study.** All oligos were synthesized by Dharmacon (Horizon). f indicates a 2'fluorine modification. * indicates a phosphorothioate modification.

| Oligo Name | Oligo Sequence | Purpose | Length (nt) |
|---|---|---|---|
| AU$^{f}$A | A.U$^{f}$.A. | Cryo-EM | 3 |
| AUA | A.U.A. | Cryo-EM | 3 |
| Fl-control | Fl.A.A.A.U. | FRET | 4 |
| TAMRA-control | A.A.TAMRA | FRET | 2 |
| Fretmer-1 | Fl.A.A.A.U.A.A.TAMRA | FRET | 6 |
| Fretmer-2 | Fl.A.A.A.A.A.A.TAMRA | FRET | 6 |
| Fretmer-3 | Fl.A.A.A.C.A.A.TAMRA | FRET | 6 |
| Fretmer-4 | Fl.A.A.C.U.A.A.TAMRA | FRET | 6 |
| Fretmer-5 | Fl.A.A.G.U.A.A.TAMRA | FRET | 6 |
| Fretmer-6 | Fl.A.A.A.U.C.A.TAMRA | FRET | 6 |
| Fretmer-7 | Fl.A.A.A.U.G.A.TAMRA | FRET | 6 |
| Fretmer-8 | Fl.A.A.U*U.A.A.TAMRA | FRET | 6 |
| Fretmer-9 | Fl.A.A.U.U.A.A.TAMRA | FRET | 6 |
| Fretmer-10 | Fl.A.A.A.U*U.A.TAMRA | FRET | 6 |
| Fretmer-11 | Fl.A.A.A.U.U.A.TAMRA | FRET | 6 |
| TRS-N | Fl.U.C.A.U.C.U.A.A.A.C.G.A.A.C.A.A.A.C.U.A.<br>A.A.A.Cy5 | Gel cleavage | 23 |
| TRS-S | Fl.A.A.C.A.A.C.U.A.A.A.C.G.A.A.C.A.A.U.Cy5 | Gel cleavage | 18 |
| Eicosamer | Fl.G.A.A.C.U.C.A.U.G.G.A.C.C.U.U.G.G.C.A.<br>G.Cy5 | Gel cleavage | 20 |
| Fl-U17 | Fl.U.U.U.U.U.U.U.U.U.U.U.U.U.U.U.U.U | Gel cleavage | 17 |
| PolyUplusNS | Fl.U.U.U.U.U.U.U.U.U.U.U.U.U.U.U.U.U.U.U.U.<br>U.U.U.U.U.U.U.U.U.U.U.U.U.G.U.C.A.U.U.<br>C.U.C.C.U.A.A.G.A.A.G.C. Cy5 | Gel cleavage | 51 |
| WTNS | G.U.C.A.U.U.C.U.C.C.U.A.A.G.A.A.G.C. Cy5 | Gel cleavage | 18 |
| PolyUplusNoUNS | Fl.U.U.U.U.U.U.U.U.U.U.U.U.U.U.U.U.U.U.U.U.<br>U.U.U.U.U.U.U.U.U.U.U.U.U.G.A.C.A.A.A.C<br>.A.C.C.A.A.A.G.A.A.G.C. Cy5 | Gel cleavage | 51 |
| NoUNS | G.A.C.A.A.A.C.A.C.C.A.A.A.G.A.A.G.C. Cy5 | Gel cleavage | 18 |
| PolyUNoYNS | Fl.U.U.U.U.U.U.U.U.U.U.U.U.U.U.U.U.U.U.U.U.<br>U.U.U.U.U.U.U.U.U.U.U.U.U.G.A.G.A.A.A.G<br>.A.G.G.A.A.A.G.A.A.G.A .Cy5 | Gel cleavage | 51 |
| NoYNS | G.A.G.A.A.A.G.A.G.G.A.A.A.G.A.A.G.A .Cy5 | Gel cleavage | 18 |

**Supplementary Table 2.** Nsp15-AUA interaction energy calculations for the hexamer system. HIE refers to histidine protonated on $N_{\varepsilon}$. HIP refers to histidine with in the doubly-protonated form, with hydrogens on both ring nitrogens (i.e. positively charged). Averaged MM/GBSA interaction energies in kcal/mol (standard deviations given in parentheses) calculated from the Nsp15-AUA hexamer assembly. The energies were calculated over all monomer conformations from the three 500 ns hexamer trajectories (50-ns segments with 50 conformations in each segment). Energies from the segments in which the RNA timer left the binding site were not included in averaging. Only the residues that had magnitudes of interaction energies larger than 0.5kcal/mol were tabulated.

| Residue | $A(B_{-2})$ | $U(B_{-1})$ | $A(B_{+1})$ |
|---|---|---|---|
| E234 | 0.00(0.00) | 0.01(0.01) | -0.71(0.80) |
| H235 (HIE) | -0.03(0.03) | -0.34(0.36) | -4.57(2.24) |
| H243 (HIE) | -0.04(0.29) | 0.00(0.02) | -0.56(0.93) |
| Q245 | -0.41(1.00) | -0.87(0.89) | -1.46(1.70) |
| G247 | -0.02(0.01) | -0.74(0.26) | -3.29(1.00) |
| G248 | -0.01(0.01) | -0.57(0.26) | -3.56(0.94) |
| H250 (HIP) | -0.05(0.03) | -5.46(1.76) | -9.61(2.26) |
| N278 | -0.01(0.02) | -1.36(1.38) | -0.02(0.02) |
| K290 | -0.41(0.67) | -7.97(3.29) | -4.82(3.79) |
| V292 | -0.15(0.33) | -1.78(0.93) | -0.10(0.06) |
| C293 | -0.03(0.04) | -1.72(0.67) | -0.16(0.11) |
| S294 | -0.00(0.01) | -2.02(1.28) | 0.09(0.15) |
| M331 | -0.56(0.56) | -0.68(0.47) | -0.15(0.10) |
| W333 | -3.20(0.95) | -1.77(0.91) | -0.61(0.67) |
| E340 | -0.59(0.75) | -0.12(0.06) | -2.45(1.36) |
| T341 | -0.51(0.43) | -0.91(0.28) | -4.90(1.26) |
| Y343 | -1.24(0.56) | -7.16(1.40) | -0.90(0.68) |
| P344 | 0.00(0.02) | -0.83(0.76) | 0.22(0.07) |
| K345 | -1.01(1.32) | -3.63(2.47) | 0.04(0.04) |

**Supplementary Table 3.** Nsp15-AUA interaction energy calculations for the monomer system. HIE refers to histidine protonated on $N_{\varepsilon}$. HIP refers to histidine with in the doubly-protonated form, with hydrogens on both ring nitrogens (i.e. positively charged). Averaged MM/GBSA interaction energies from an Nsp15-AUA monomer system. The values were calculated over three 500 ns independent trajectories (50 ns segments with 50 conformations in each segment) and the segments in which the RNA trimer left the binding site were not included in the averaging. Only the residues that had magnitudes of interaction energies larger than 0.5kcal/mol were tabulated.

| Residue | A($B_{-2}$) | U($B_{-1}$) | A($B_{+1}$) |
|---|---|---|---|
| H235 (HIE) | -0.07(0.11) | -0.30(0.26) | -2.56(1.41) |
| Q245 | -0.12(0.44) | -0.50(0.45) | -3.47(1.52) |
| L246 | -0.02(0.01) | 0.11(0.17) | -0.95(1.06) |
| G247 | -0.03(0.02) | -0.85(0.18) | -4.40(0.83) |
| G248 | -0.03(0.02) | -0.58(0.21) | -3.81(0.79) |
| H250 (HIP) | -0.19(0.09) | -7.99(2.01) | -10.49(1.90) |
| N278 | -0.50(0.51) | -1.27(1.25) | -0.04(0.02) |
| K290 | -0.45(0.51) | -10.11(3.39) | -13.01(3.76) |
| V292 | -1.13(0.65) | -2.36(0.95) | -0.26(0.14) |
| C293 | -0.43(0.35) | -2.30(0.65) | -0.22(0.12) |
| S294 | -0.21(0.33) | -3.05(1.46) | 0.15(0.16) |
| W333 | -3.22(1.14) | -0.31(0.49) | -0.20(0.49) |
| E340 | -1.26(1.08) | -0.09(0.04) | -0.47(0.39) |
| T341 | -0.94(0.38) | -0.58(0.26) | -3.77(1.39) |
| Y343 | -0.89(0.68) | -5.86(1.44) | -0.41(0.54) |
| P344 | 0.01(0.03) | -0.59(0.45) | 0.24(0.06) |
| K345 | -1.15(1.34) | -3.26(3.27) | 0.03(0.06) |

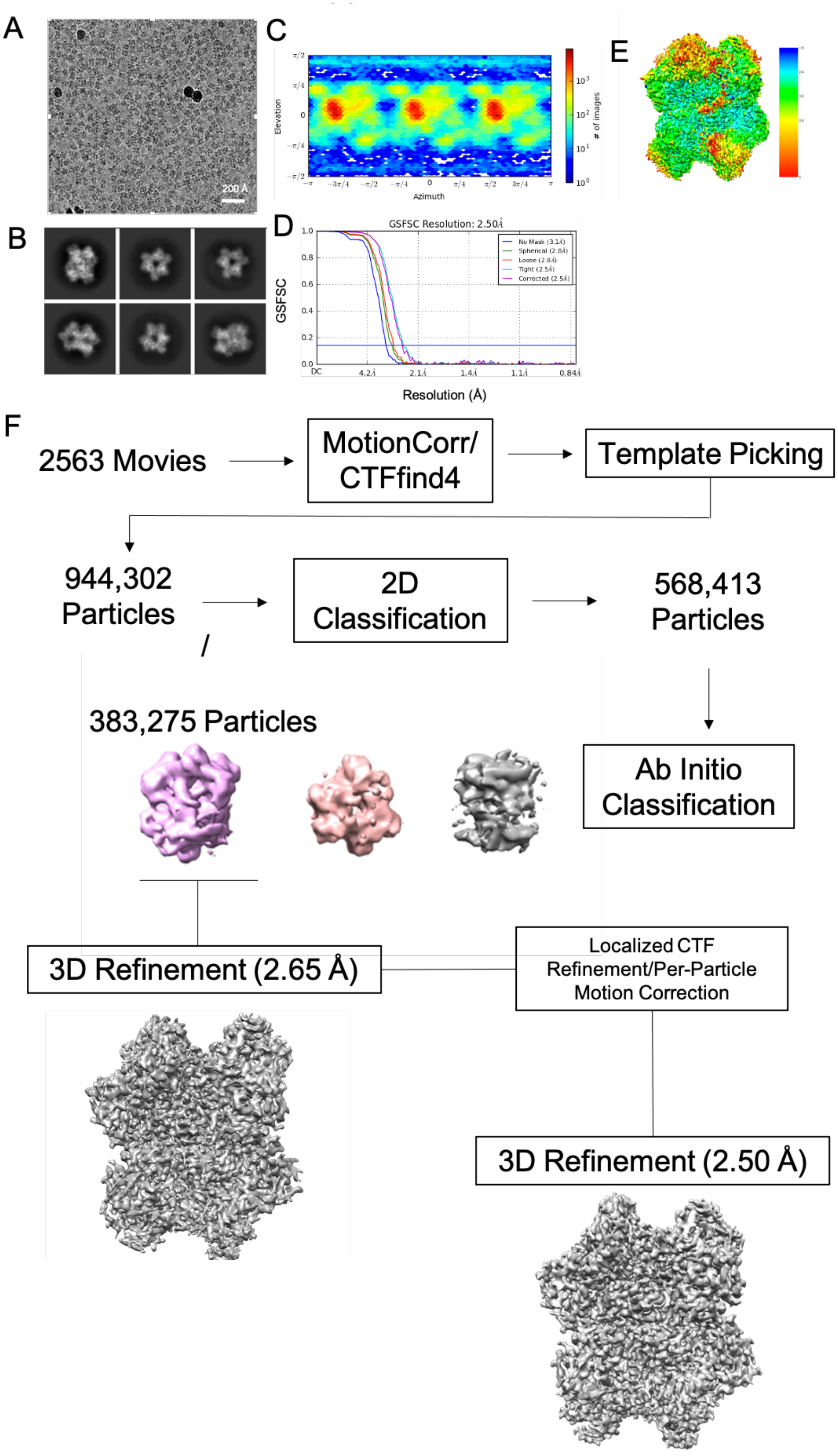
A
200 Å
B
C
Elevation
Azimuth
### of images
D
GSFSC Resolution: 2.50Å
GSFSC
Resolution (Å)
No Mask (3.1Å)
Spherical (2.8Å)
Loose (2.8Å)
Tight (2.5Å)
Corrected (2.5Å)
E
F
2563 Movies
MotionCorr/ CTFfind4
Template Picking
944,302 Particles
2D Classification
568,413 Particles
383,275 Particles
Ab Initio Classification
3D Refinement (2.65 Å)
Localized CTF Refinement/Per-Particle Motion Correction
3D Refinement (2.50 Å)

**Supplementary Fig. 1.** Cryo-EM processing workflow for the pre-cleavage state. (**A**) A representative micrograph of WT-Nsp15 with excess $AU^{f}A$ in vitreous ice and (**B**) selected 2D classes generated from 2563 movies collected from an UltrAuFoil R1.2/1.3 300 mesh grid. (**C**) Angular distribution of pre-cleavage Nsp15 particles. (**D**) Fourier shell correlation (FSC) curve for the pre-cleavage Nsp15 reconstruction. The overall resolution is 2.50 Å according to the FSC 0.143 criteria (1,2). (1,2) (**E**) Cryo-EM reconstruction of the pre-cleavage state colored based on local resolution calculated using cryoSparc v2 (3). (**F**) Cryo-EM processing workflow. Picked particles (944,302) were subjected to 2D classification, 3D classification, and refinement prior to local refinement of a single Nsp15 protomer in cryoSPARC v2 (3).

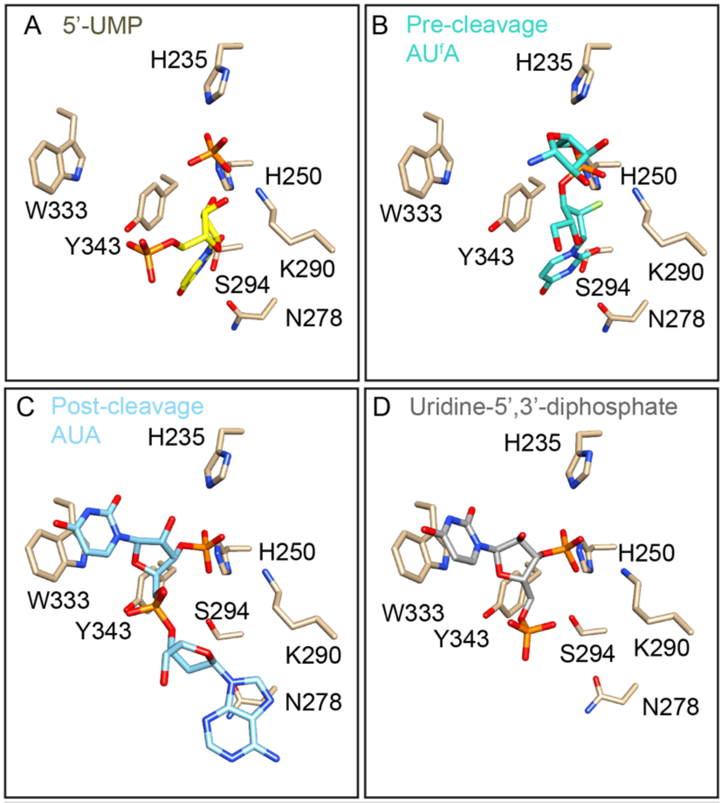


**Supplementary Fig. 2. Nsp15 structure comparison.** In all panels, Nsp15 active site residues are shown in side chain representation (tan). (**A**) Nsp15 with 5′-UMP (yellow, PDB: 7K0R, (4)). (**B**) Nsp15 with AU$^{f}$A (turquoise, pre-cleavage structure, this work). (**C**) Nsp15 with AUA (sky blue, post-cleavage structure, this work). (**D**) Nsp15 with uridine-5′,3′-diphosphate (grey, PDB:7K1O).

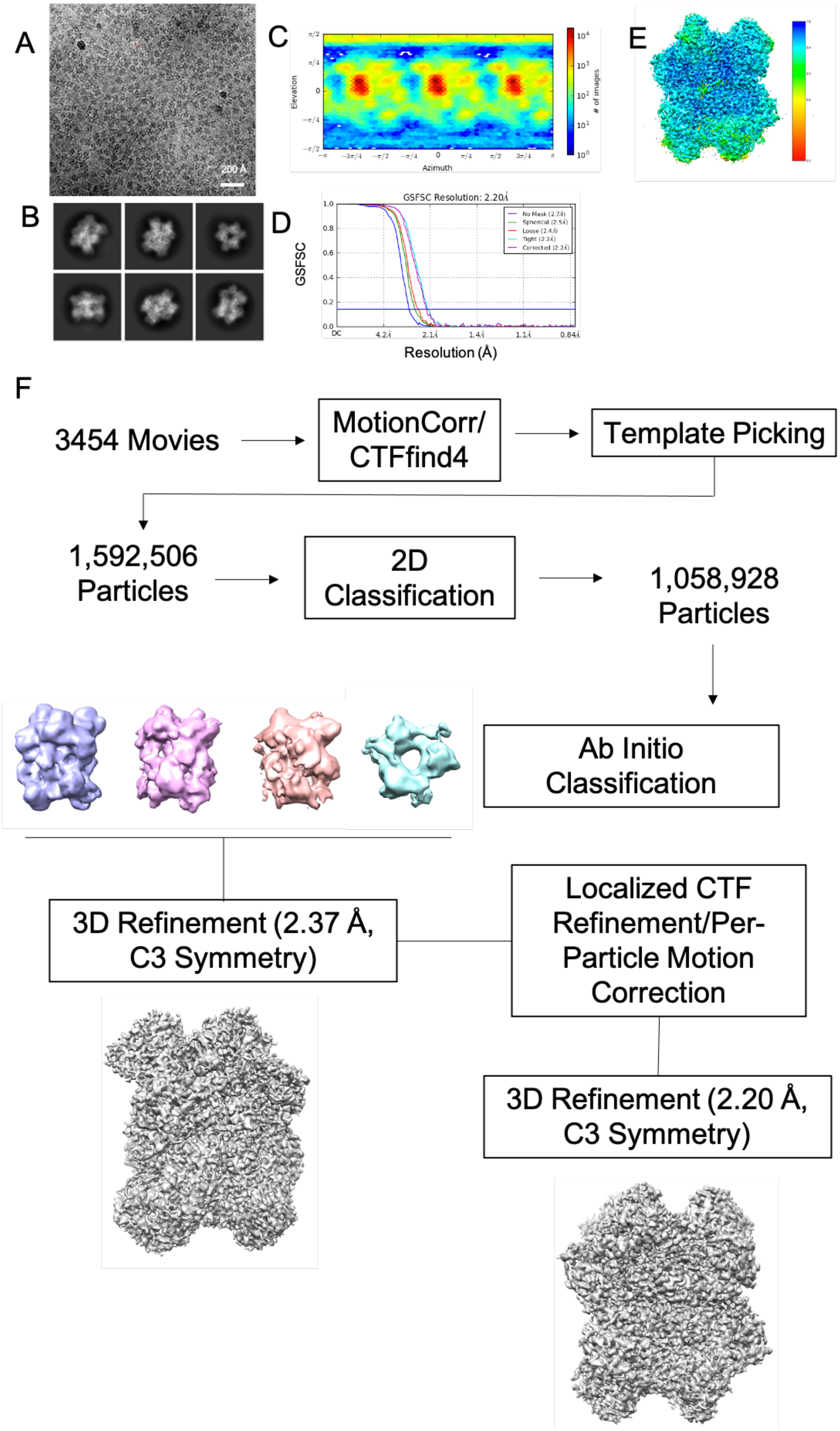
A
200 Å
B
C
Elevation
Azimuth
### of images
D
GSFSC Resolution: 2.20Å
GSFSC
No Mask (2.7Å)
Spherical (2.5Å)
Loose (2.4Å)
Tight (2.2Å)
Corrected (2.2Å)
DC
4.2Å
2.1Å
1.4Å
1.1Å
0.84Å
Resolution (Å)
E
F
3454 Movies
MotionCorr/ CTFfind4
Template Picking
1,592,506 Particles
2D Classification
1,058,928 Particles
Ab Initio Classification
3D Refinement (2.37 Å, C3 Symmetry)
Localized CTF Refinement/Per-Particle Motion Correction
3D Refinement (2.20 Å, C3 Symmetry)

**Supplementary Fig. 3. Cryo-EM processing workflow for the post-cleavage state.** Cryo-EM processing workflow for the post-cleavage state. (**A**) A representative micrograph of WT-Nsp15 with excess AUA in vitreous ice and (**B**) selected 2D classes generated from 3454 movies collected from an UltrAuFoil R1.2/1.3 300 mesh grid. (**C**) Angular distribution of post-cleavage Nsp15 particles. (**D**) Fourier shell correlation (FSC) curve for the post-cleavage Nsp15 reconstruction. The overall resolution is 3.38 Å according to the FSC 0.143 criteria (1,2). (**E**) Cryo-EM reconstruction of the post-cleavage state colored based on local resolution calculated using cryoSparc v2 (3). (**F**) Cryo-EM processing workflow. Picked particles (1,592,506) were subjected to 2D classification, 3D classification, and refinement prior to local refinement of a single Nsp15 protomer in cryoSPARC v2 (3).

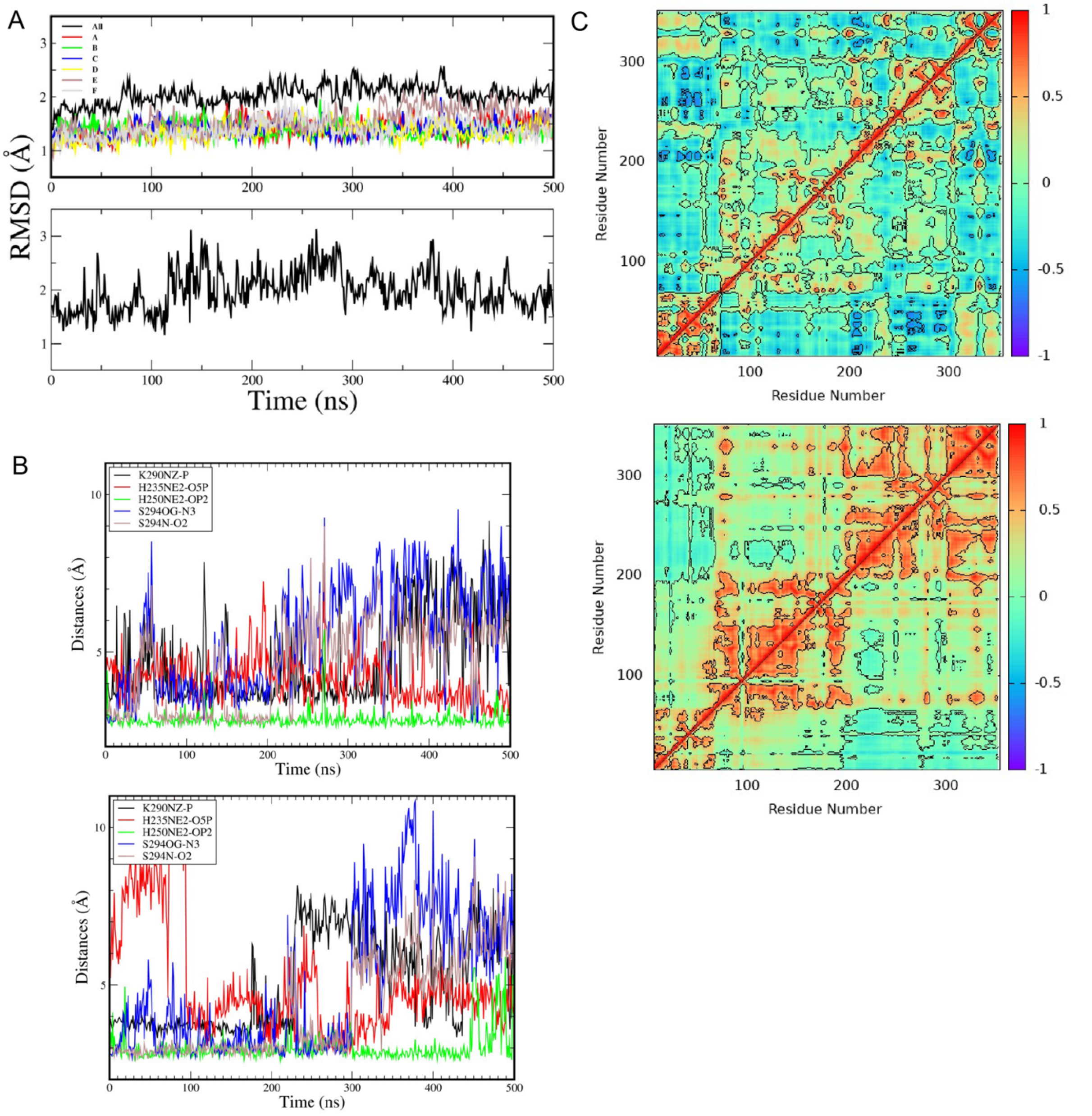


**Supplementary Fig. 4.** Molecular dynamics simulations of RNA binding. (**A**) Root mean squared deviations (RMSD) calculated with respect to the starting conformations for a representative hexameric (top) and a monomeric (bottom) system from three samples of each system. Stability of the solvated monomers and hexamers during 0.5 µs dynamics was established using these RMSD plots. (**B**) Various distances between Nsp15 and the trinucleotide atoms calculated from the molecular dynamics trajectories of the hexamer (top) and the monomer (bottom) were used to select whether the trinucleotide was in a bound conformation. (**C**) Dynamic cross-correlation matrix analysis helped visualize highly correlated dynamics among the monomer subdomains in a hexamer trajectory (bottom) as oppose to less correlated dynamics in a monomer trajectory (top). The monomer subdomains are the small N-terminal domain (ND: residues 1-64), the middle domain (MD: 65-183), followed by the conserved endoU domain (residues 184-347) containing the nuclease active site. For both (**B**) and (**C**) a representative trajectory from the three samples of the monomer systems was selected as well as a representative protomer from a hexamer trajectory out of the 18 available protomers within the hexameric units in the three trajectories.

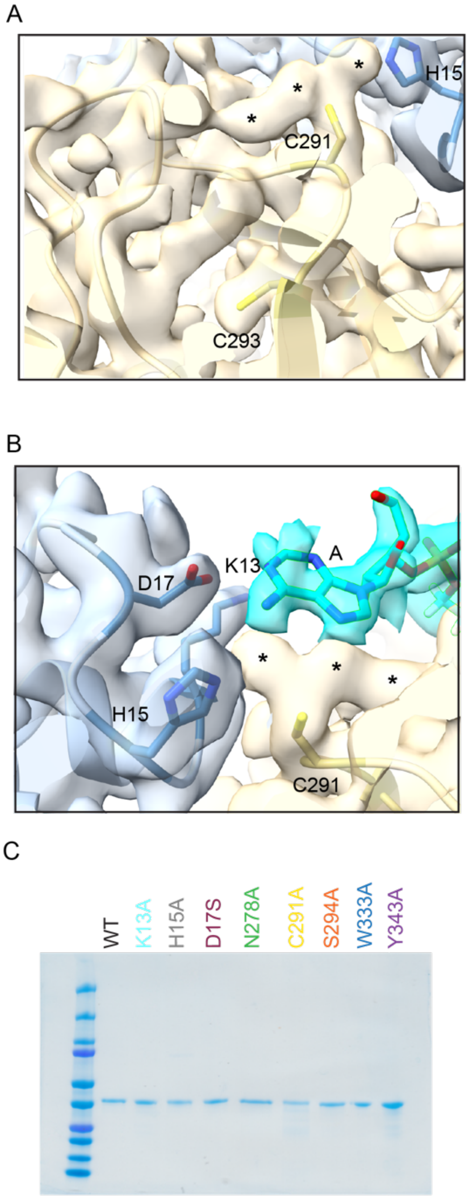


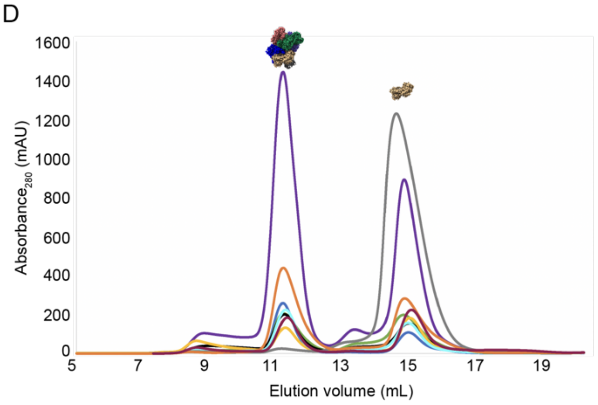


E

| Construct | Ratio (Hexamer/Monomer) |
|---|---|
| WT | 1.11 |
| K13A | 1.21 |
| H15A | 0.02 |
| D17S | 0.79 |
| N278A | 0.87 |
| C291A | 0.71 |
| S294A | 1.41 |
| W333A | 2.04 |
| Y343A | 1.56 |

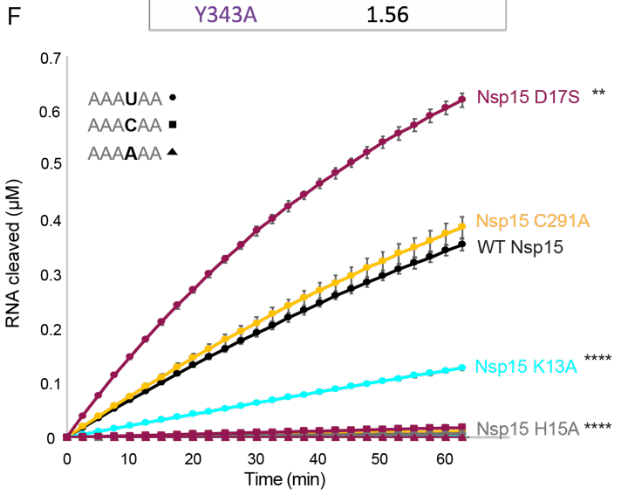


**Supplementary Fig. 5.** N-terminal domain active site contributions. (**A-B**) The post-cleavage map and model is shown zoomed into the area where the $B_{-2}$ adenine interacts. The neighboring chain is colored in blue and the RNA is colored in teal. The map was colored by chain using the distance cut off of 2.5Å. The resolution of the map results in well-ordered side chain density, and lead to the observation of extra density (*) extending from C291. (**A**) No extra density is seen around C293 compared to C291. (**B**) The image is rotated 180˚ from (A) and K13, H15, and D17 are shown along with the $B_{-2}$ adenine density. (**C**) SDS-PAGE protein gel of all constructs used in this work, illustrating pure protein. The constructs are colored according to the chromatography and activity curves. (**D**) Overlaid S200 size exclusion chromatography traces are shown, with the hexamer peak eluting before the monomer peak. Traces are labeled in the corresponding color shown on the SDS-PAGE gel. (**E, F**) FRET reaction curves show C291 is not critical, as mutation to alanine does not affect cleavage. Nsp15 variants (2.5 nm) were incubated with RNA substrates (0.8 µM) at RT for 60 minutes, with fluorescence read every 2.5 minutes. The average of a representative technical triplicate is plotted with standard deviation error bars. At least two biological replicates were performed for each mutant; Dunnett's T3 multiple corrections test was performed using Prism for pairwise comparisons. The p-values are ** $p = 0.0038$, **** $p < 0.0001$. Three different substrates were tested for each mutant: AAA**U**AA (circles), AAA**A**AA (squares), and AAA**C**AA (triangles). Each mutant is represented by a different color: WT Nsp15 (black), Nsp15 K13A (cyan), Nsp15 H15A (grey), Nsp15 D17S (maroon), Nsp15 C291A (yellow).

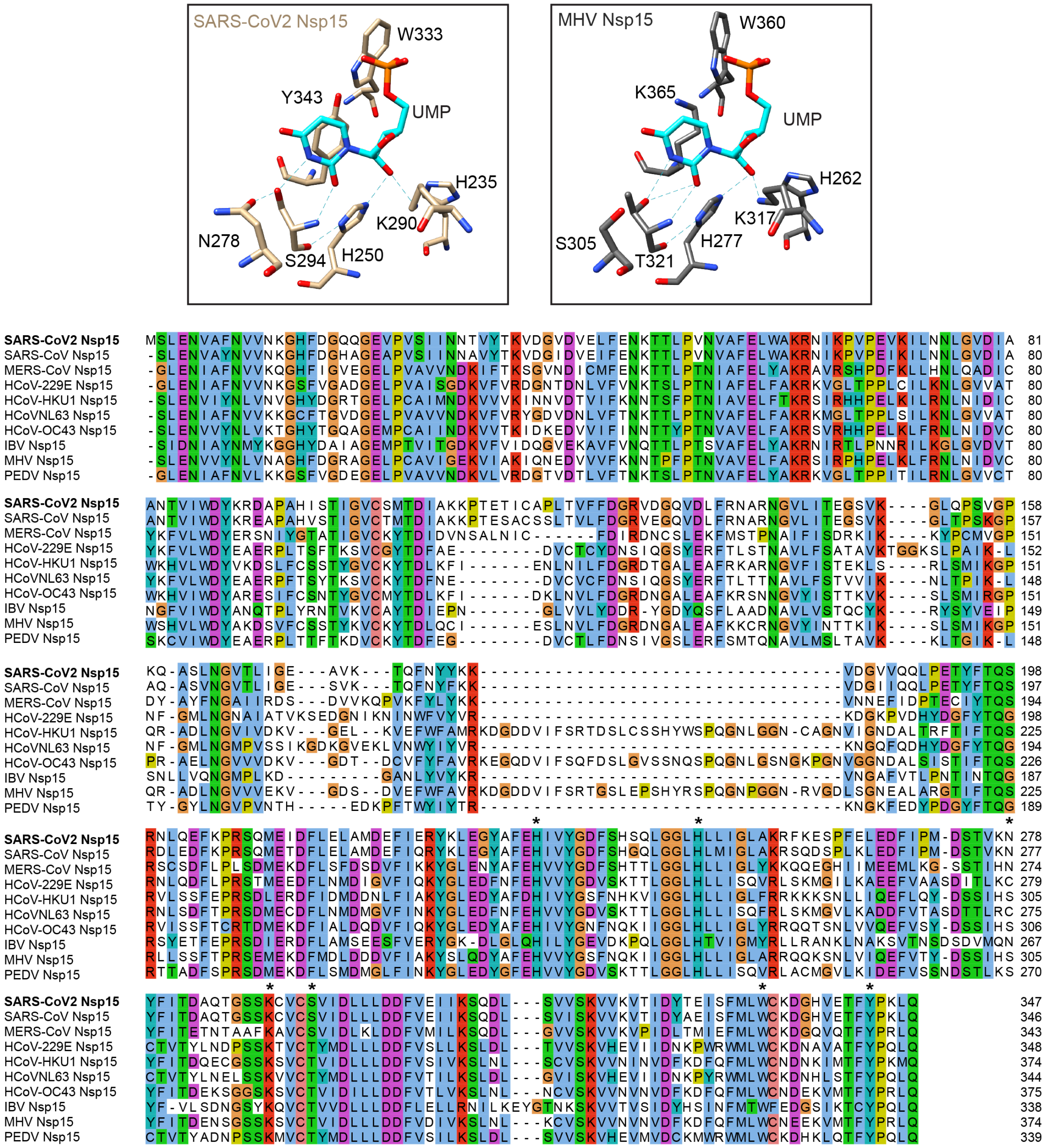


**Supplementary Fig. 6.** Comparison with other coronavirus Nsp15 sequences. Top: Crystal structure alignment of SARS-CoV-2 Nsp15 (PDB: 6X4I) with 3'UMP bound and MHV Nsp15 (PDB: 2GTH). Ligand is colored in teal, SARS-CoV-2 Nsp15 in tan, and MHV Nsp15 in gray. Hydrogen bonds are depicted by green lines. Selected active site residues are labeled. Bottom: Full sequence alignment for Nsp15 protein sequences from the human coronaviruses as well as three major animal coronaviruses. The residues of interest are marked with an asterisk. Alignment was created in Jalview (5).

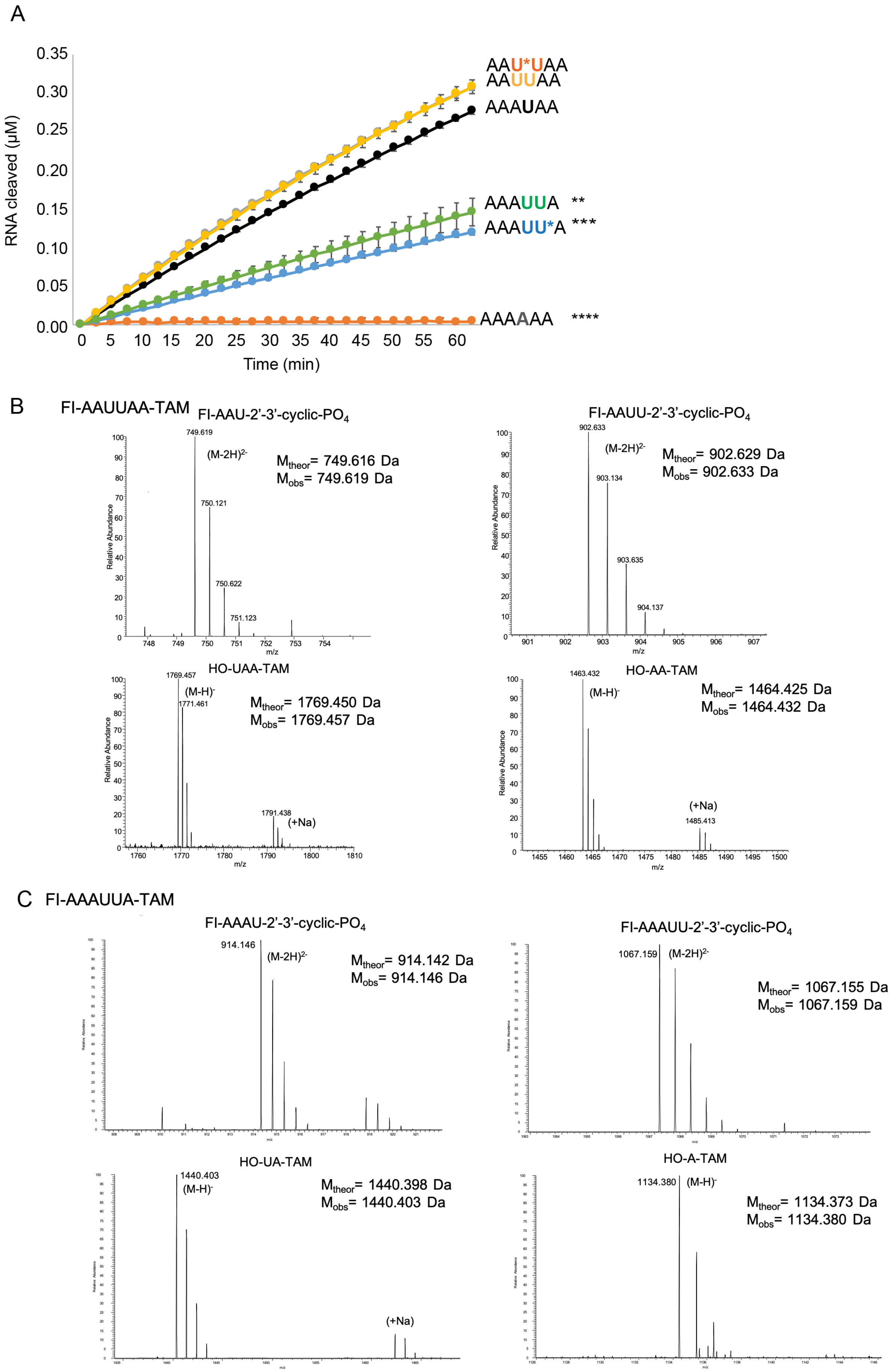
A
RNA cleaved (μM)
Time (min)
AAU*UAA
AAUUAA
AAAUAA
AAAUUA **
AAAUU*A ***
AAAAAA ****
B
Fl-AAUUAA-TAM
Fl-AAU-2'-3'-cyclic-PO4
(M-2H)2-
Mtheor= 749.616 Da
Mobs= 749.619 Da
Fl-AAUU-2'-3'-cyclic-PO4
(M-2H)2-
Mtheor= 902.629 Da
Mobs= 902.633 Da
HO-UAA-TAM
(M-H)-
Mtheor= 1769.450 Da
Mobs= 1769.457 Da
(+Na)
HO-AA-TAM
(M-H)-
Mtheor= 1464.425 Da
Mobs= 1464.432 Da
(+Na)
C
Fl-AAAUUA-TAM
Fl-AAAU-2'-3'-cyclic-PO4
(M-2H)2-
Mtheor= 914.142 Da
Mobs= 914.146 Da
Fl-AAAUU-2'-3'-cyclic-PO4
(M-2H)2-
Mtheor= 1067.155 Da
Mobs= 1067.159 Da
HO-UA-TAM
(M-H)-
Mtheor= 1440.398 Da
Mobs= 1440.403 Da
(+Na)
HO-A-TAM
(M-H)-
Mtheor= 1134.373 Da
Mobs= 1134.380 Da
Relative Abundance
m/z

**Supplementary Fig. 7.** Extended FRET assay and mass spectrometry analysis. (**A**) FRET time course data for WT Nsp15 (2.5 nm) incubated with indicated RNA substrates (0.8 μM) at RT for 60 minutes, with fluorescence read every 2.5 minutes. * indicates a phosphorothioate modification, which is a non-hydrolyzable bond. The average of a representative technical triplicate is plotted with standard deviation error bars; Dunnett's T3 multiple corrections test was performed using Prism for pairwise comparisons. The p-values are ** $p = 0.0028$, *** $p = 0.0002$, **** $p < 0.0001$. (**B-C**) LC-MS was performed on the cleavage products resulting from NSP15 treatment of FI-AAUUAA-TAM (**B**) and FI-AAAUUA-TAM (**C**). MS data shows that cleavage was readily detected between the uridines (left sides of panels B and C) as well as 3' of the second uridine (right sides of panels B and C).

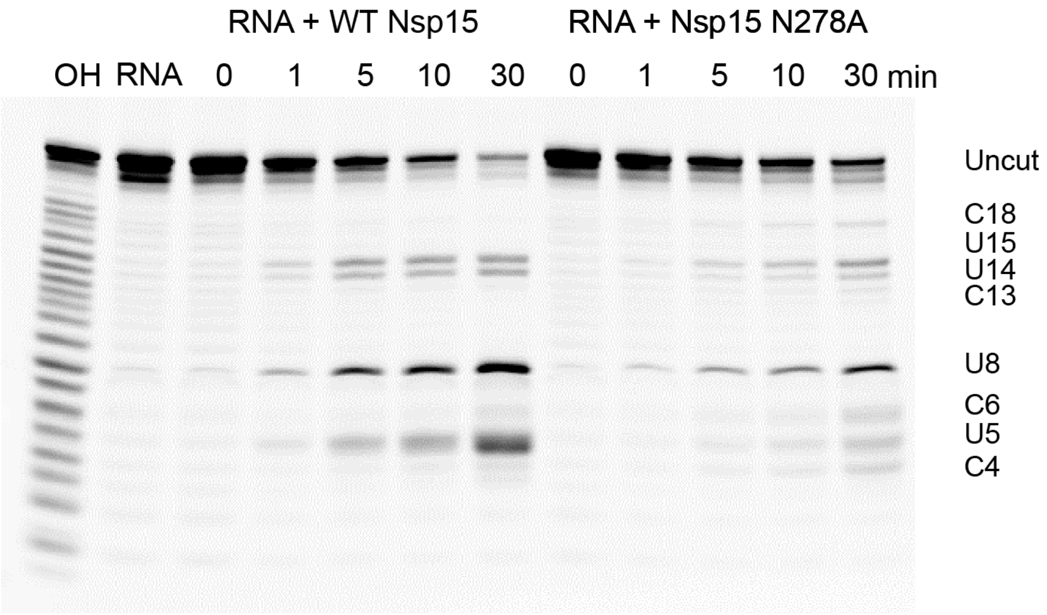


**Fl**.G.A.A.$C_4$.$U_5$.$C_6$.A.$U_8$.G.G.A.$C_{12}$.$C_{13}$.$U_{14}$.$U_{15}$.G.G.$C_{18}$.A.G.Cy5

**Supplementary Fig. 8.** Nsp15 prefers purines 3′ of uridines in longer substrates. A 20-mer oligo (500 nM) with multiple uridines was incubated with Nsp15 (50 nM, WT, left or N278A, right) for a 30-minute time course at room temperature. The products were resolved with a 20% acrylamide gel and the image for the 5′-FI labeled products is shown. Cy5 label anomalous migration affected interpretation of 3′-products. The products are labeled by nucleotide number. Each reaction was performed three times, including two biological replicates.

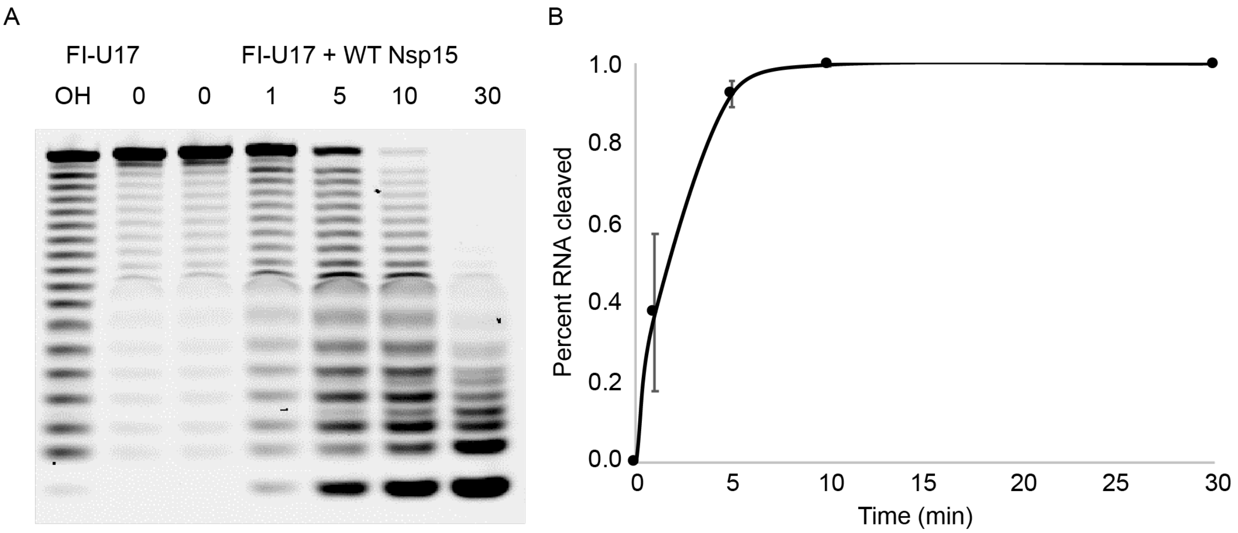


**Supplementary Fig. 9.** Nsp15 efficiently cleaves polyU RNA. (**A**) Gel-based cleavage assay with FI-U17 substrate (500 nM) and WT Nsp15 (50 nM) over a 30-time course reaction at room temperature. Alkaline hydrolysis for 15 min at 90˚ was used to create a ladder. (**B**) The disappearance of the uncut band was quantified from 3 replicates and graphed as the percent RNA cleaved (the inverse of the disappearance) normalized to time 0. The reaction was performed three times, with two biological replicates.

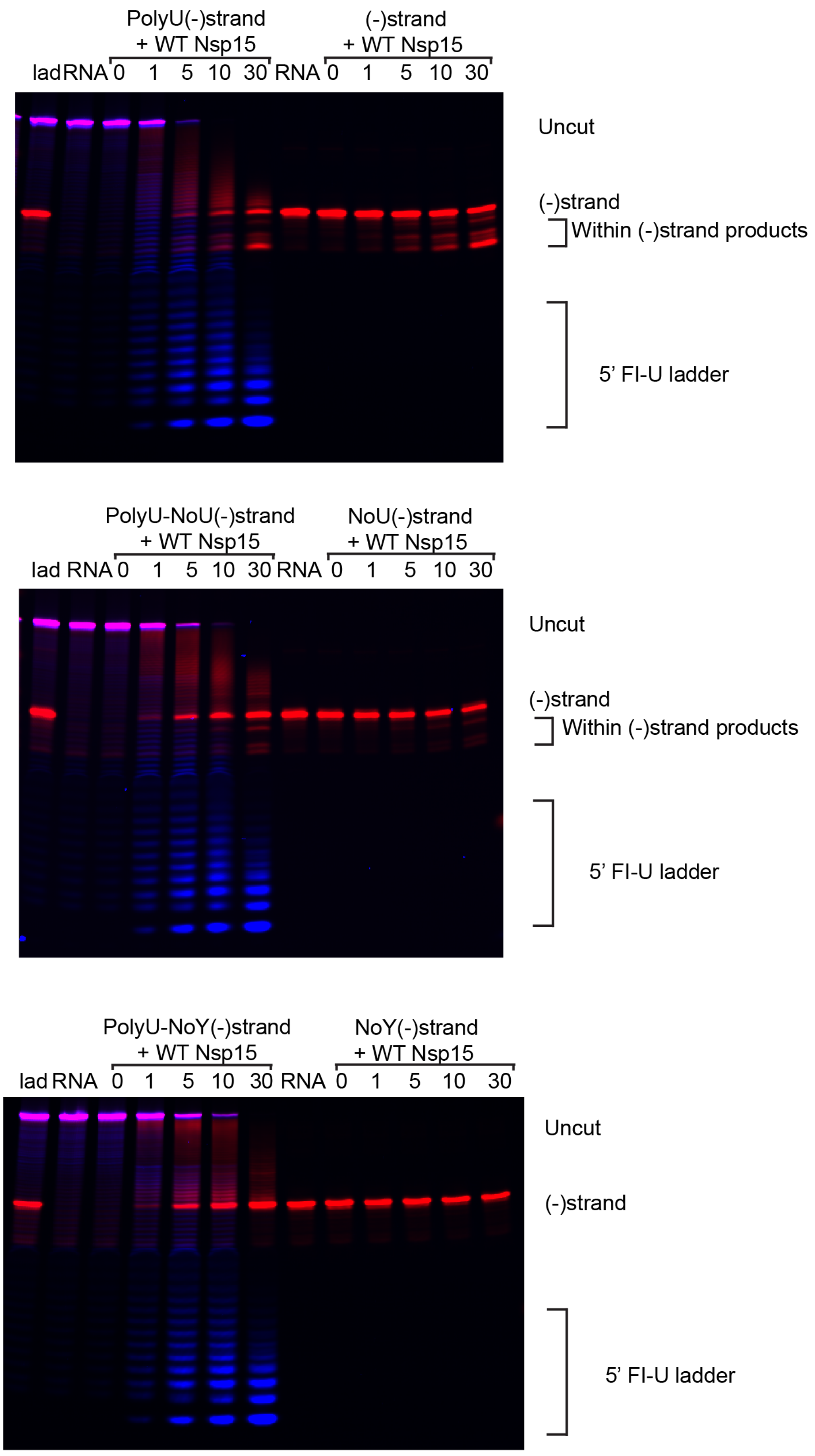
PolyU(-)strand
+ WT Nsp15
(-)strand
+ WT Nsp15
lad RNA 0 1 5 10 30 RNA 0 1 5 10 30
Uncut
(-)strand
Within (-)strand products
5’ FI-U ladder
PolyU-NoU(-)strand
+ WT Nsp15
NoU(-)strand
+ WT Nsp15
lad RNA 0 1 5 10 30 RNA 0 1 5 10 30
Uncut
(-)strand
Within (-)strand products
5’ FI-U ladder
PolyU-NoY(-)strand
+ WT Nsp15
NoY(-)strand
+ WT Nsp15
lad RNA 0 1 5 10 30 RNA 0 1 5 10 30
Uncut
(-)strand
5’ FI-U ladder

**Supplementary Fig. 10.** Individual gels for each negative strand cleavage reaction. The polyU substrates are labeled with 5′-Fl (blue) and 3′-Cy5 (red) while the negative strand substrates only have a 3′-Cy5 (red) label. 500 nM substrate was incubated with 50 nM WT Nsp15 at room temperature for 30 minutes. Products were resolved on a 15% acrylamide gel. Each reaction was performed three times, including two biological replicates.
